# Cardiac function modulates endocardial cell dynamics to shape the cardiac outflow tract

**DOI:** 10.1101/787358

**Authors:** Pragya Sidhwani, Giulia L.M. Boezio, Hongbo Yang, Neil C. Chi, Beth L. Roman, Didier Y.R. Stainier, Deborah Yelon

**Affiliations:** Division of Biological Sciences, University of California, San Diego, La Jolla, CA, 92093, USA; Max Planck Institute for Heart and Lung Research, Department of Developmental Genetics, 61231 Bad Nauheim, Germany; Division of Cardiovascular Medicine, Department of Medicine, University of California, San Diego, La Jolla, CA, 92093, USA; Department of Human Genetics, Graduate School of Public Health, and Heart, Lung, Blood and Vascular Medicine Institute, University of Pittsburgh, Pittsburgh, PA, 15261, USA

## Abstract

Physical forces are important participants in the cellular dynamics that shape developing organs. During heart formation, for example, contractility and blood flow generate biomechanical cues that influence patterns of cell behavior. Here, we address the interplay between function and form during the assembly of the cardiac outflow tract (OFT), a crucial connection between the heart and vasculature that develops while circulation is underway. In zebrafish, we find that the OFT expands via accrual of both endocardial and myocardial cells. However, when cardiac function is disrupted, OFT endocardial growth ceases, accompanied by reduced proliferation and reduced addition of cells from adjacent vessels. The TGFβ receptor Acvrl1 is required for addition of endocardial cells, but not for their proliferation, indicating distinct regulation of these essential cell behaviors. Together, our results suggest that cardiac function modulates OFT morphogenesis by triggering endocardial cell accumulation that induces OFT lumen expansion and shapes OFT dimensions.

## INTRODUCTION

Organs are sculpted in a dynamic physical environment, where they are pulled, squeezed and twisted in a spatiotemporally heterogeneous manner. Such anisotropic mechanical influences are indispensable for proper development: they activate specific signaling cascades to drive essential cellular decisions regarding growth and rearrangement (Eder et al., 2017; Heisenberg & Bellaiche, 2013). In this way, physical influences forge an organ’s characteristic contours and proportions, which ultimately facilitate its function. Despite their importance, however, the precise mechanisms that tune organ shape in response to forces remain elusive in many contexts.

The embryonic heart serves as an important example of how biomechanical cues enforce organ form (Collins & Stainier, 2016; Lindsey et al., 2014; Sidhwani & Yelon, 2019). As blood flows through the cardiac chambers, it sloshes along the grooves and gorges created by the inner endocardial layer of the heart. The resulting perturbations in fluid parameters, as well as contractility intrinsic to the outer myocardial layer, can jointly modulate cell behaviors to create a heart of the appropriate dimensions. In the zebrafish heart, for example, blood flow promotes regional patterns of cellular expansion and myofibril maturation, while contractility restricts myocardial cell size, as the characteristic curvatures of the ventricular chamber emerge (Auman et al., 2007; Lin et al., 2012). This myocardial morphogenesis is spatiotemporally coordinated with that of the underlying ventricular endocardium, where blood flow instigates proliferation and cellular hypertrophy (Dietrich et al., 2014). In a similar timeframe, flow-induced expression of the TGFβ type I receptor Acvrl1 in arterial vessels facilitates migration of endothelial cells into the heart, resulting in the restriction of the diameter of the distal cranial vessels (Corti et al., 2011; Rochon et al., 2016). Flow-related parameters also activate the calcium channels Trpp2 and Trpv4 to upregulate expression of the transcription factor gene *klf2a*, which remodels the endocardial lining of the atrioventricular canal in a *wnt9b*-dependent manner (Galvez-Santisteban et al., 2019; Goddard et al., 2017; Heckel et al., 2015). Thus, cardiac function impacts multiple tissues – myocardium, endocardium, and endothelium – during the synchronous development of the cardiovascular system.

The cardiac outflow tract (OFT), an essential tubular junction where the heart meets the vessels, plays a pivotal role in the collaboration between cardiac and vascular morphogenesis. Moreover, errors in OFT formation cause a variety of cardiac birth defects: as many as 30% of all cases of congenital heart disease feature OFT malformations (Neeb et al., 2013; Pierpont et al., 2018). In all vertebrates, OFT development begins with the assembly of a small tube of myocardium, derived from late-differentiating second heart field (SHF) cells that append to the arterial pole of the embryonic ventricle (Kelly, 2012; Knight & Yelon, 2016; Poelmann & Gittenberger-de Groot, 2019). The precise dimensions of this tube are essential, since this myocardial investment provides the foundation for subsequent remodeling events, including septation, cushion formation, and rotation, that ultimately create a mature OFT aligned with the arterial vasculature. Defects in these later processes can originate with flaws in the initial construction of the OFT tube, highlighting the importance of understanding the regulatory mechanisms that carefully calibrate its dimensions.

Previous studies have provided extensive insights into the pathways regulating SHF proliferation, epithelialization and differentiation, which ensure an adequate supply of myocardial cells for the assembly of the initial OFT (Dyer & Kirby, 2009; Kelly, 2012; Knight & Yelon, 2016; Paffett-Lugassy et al., 2017). The Fgf and canonical Wnt pathways, for example, regulate proliferation of SHF progenitor cells (Francou et al., 2013; Rochais et al., 2009). Tbx1-dependent epithelial characteristics and protrusive activity of SHF progenitors appear to be important for their proper proliferation and differentiation (Cortes et al., 2018; Francou et al., 2014; Nevis et al., 2013), and recent analyses in zebrafish also suggest that SHF-derived cells constitute an epithelial sheath that wraps around the arterial end of the endocardium (Felker et al., 2018). Subsequently, the AP-1 transcription factor Fosl2 contributes to control of the timing of SHF differentiation (Jahangiri et al., 2016). Beyond our understanding of SHF progenitor dynamics, however, not much is known about how myocardial and endocardial cells organize into an appropriately shaped OFT. Specifically, what are the cellular and molecular mechanisms that build a tract with the proper dimensions, and how do they modulate structural changes in response to increasing functional demands?

Since the OFT forms in the context of active blood flow and myocardial contractility, physical forces associated with cardiac function could influence the initial steps of establishing OFT morphology. Indeed, subsequent phases of OFT morphogenesis, such as endocardial cushion remodeling and valve formation within the mature OFT, have been shown to depend on cardiac function. For example, flow stimulates the deposition of extracellular matrix proteins during valve formation in chick OFT explants (Biechler et al., 2014), consistent with earlier work in chick in which surgical obstruction of blood flow led to OFT valve anomalies (Hogers et al., 1999). Additionally, the mechanosensitive channel Piezo1 influences OFT valve development in zebrafish through its regulation of Klf2 and Notch activity in the endocardium (Duchemin et al., 2019). These studies demonstrate the sensitivity of late stages of OFT development to biomechanical influences; however, a role for cardiac function during the initial phases of OFT construction has not been evaluated.

Here, we exploit the utility of zebrafish genetics and high-resolution morphometrics to uncover function-induced mechanisms governing the cellular underpinnings of OFT shape. During the initial process of OFT assembly, we find that expansion of the OFT involves an increase in endocardial cell number, derived from two sources: endocardial proliferation and addition of endothelial cells from the neighboring aortic arches. Loss of cardiac function leads to a collapse of the OFT endocardium and inhibits endocardial cell accrual by interfering with both proliferation and addition of cells. Importantly, we find that loss of Acvrl1 function interferes with endothelial cell addition without mitigating proliferation, in a manner that is similar to mutants in which atrial function is disrupted, presumably altering the patterns of blood flow into the ventricle and OFT. Taken together, our studies suggest a model in which cardiac contractility and blood flow concertedly trigger endocardial proliferation and Acvrl1-dependent endothelial addition in order to construct an appropriately shaped endocardial scaffold that myocardial cells encase to mold a bilayered OFT. This impact of cardiac function on OFT morphology has meaningful implications for the causes of congenital heart diseases, as well as the defects associated with hereditary hemorrhagic telangiectasia type 2 (HHT2), which is caused by mutations in *ACVRL1* (Letteboer et al., 2006; Roman & Hinck, 2017).

## RESULTS

### Cellular accrual accompanies outflow tract assembly

In zebrafish, OFT assembly initiates around 24 hours post fertilization (hpf), when SHF progenitor cells begin differentiating into myocardial cells that surround an extension of the endocardium to construct a bilayered cylinder (de Pater et al., 2009; Lazic & Scott, 2011; Zhou et al., 2011). By 48 hpf, the OFT myocardium has formed a distinct structure that is separated from the characteristic curvatures of the ventricle by a defined morphological constriction (Fig. 1A,B). Subsequently, a collar of SHF-derived smooth muscle cells appends to this myocardial scaffold by 72 hpf (Grimes & Kirby, 2009; Hami et al., 2011), and the endocardial cells at the base of the OFT form cushions around 72 hpf that remodel into valve leaflets by 120 hpf (Duchemin et al., 2019).

**Figure 1.**
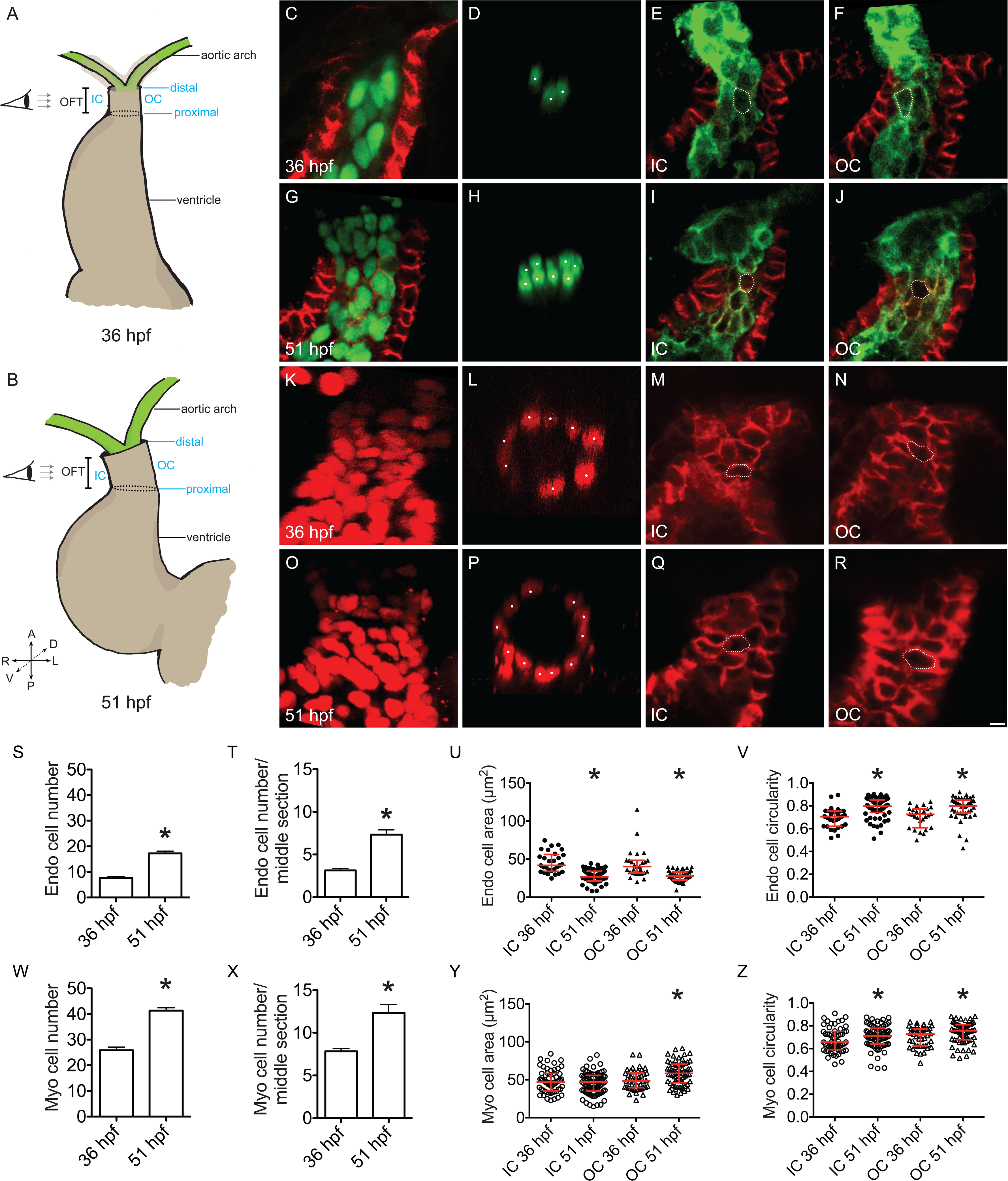
Endocardial and myocardial cells accumulate as the OFT develops. **(A,B)** Cartoons illustrate frontal views of the wild-type heart at 36 (A) and 51 hpf (B) and highlight the IC, OC, and proximal and distal boundaries of the OFT. Eye and arrows indicate the angle of imaging used to visualize the OFT. **(C,D,G,H)** Immunofluorescence indicates localization of Alcama (red), which outlines myocardial cells, and expression of *Tg(fli1a:negfp)* (green), which labels endocardial nuclei. Lateral slices (C,G) and middle cross-sections (D,H) through the OFT demonstrate that the number of OFT endocardial cells increases between 36 (C,D) and 51 hpf (G,H), as quantified in (S,T) (n=12, 9 embryos analyzed in total, over the course of 3 experimental replicates (N=3; see Materials and Methods for details regarding replicates)). Asterisks mark significant differences from 36 hpf (p<0.0001, Student’s T test). (D,H) White dots indicate GFP+ nuclei. **(E,F,I,J)** Immunofluorescence marks Alcama (red) and expression of *Tg*(*kdrl*:*HsHRAS*- *mCherry*) (green), which outlines endocardial cells. Partial reconstructions of the IC (E,I) and OC (F,J) endocardial walls demonstrate that endocardial cells get smaller (U) and rounder (V) over time (n=7, 9 embryos; 29, 59, 27, 43 cells; N=3; asterisks mark significant differences from same wall at 36 hpf, p<0.001, Mann-Whitney U test). White outlines mark representative cells. **(K,L,O,P)** Three-dimensional reconstructions (K,O) and middle cross-sections (L,P) of the OFT, with immunofluorescence marking expression of *Tg*(*myl7*:*H2A*-*mCherry*), which labels myocardial nuclei, indicate an increase in myocardial cell number between 36 (K,L) and 51 hpf (O,P), as quantified in (W,X) (n=6, 6; N=1; asterisks mark significant differences from 36 hpf, p<0.01, Student’s T test). (L,P) White dots indicate mCherry+ nuclei. **(M,N,Q,R)** Partial reconstructions of the OFT myocardial walls at 36 (M,N) and 51 hpf (Q,R), marked by immunofluorescence for Alcama, indicate trends toward more circular shape of OFT myocardial cells (Z) and enlargement of OC cells (Y) over time (n=8, 8 embryos; 57, 79, 45, 64 cells; N=2; asterisks mark significant differences from same wall at 36 hpf, p<0.05, Mann-Whitney U test). White outlines mark representative cells. **(S,T,W,X)** Bar graphs indicate mean and SEM. **(U,V,Y,Z)** Red lines represent median and interquartile range. Scale bar: 5 µm.

Although the earliest phase of OFT assembly, between 24 and 48 hpf, is an essential prerequisite for the later phases of OFT morphogenesis, we do not yet have a high-resolution understanding of the cell behaviors that underlie the dimensions of the initial OFT tube. We therefore chose to examine multiple cellular characteristics of the OFT as it expands between two stages: 36 hpf, when differentiation of SHF cells into OFT myocardium is underway, and 51 hpf, when the full span of OFT myocardium has been established (Guner-Ataman et al., 2013; Lazic & Scott, 2011; Zhou et al., 2011). Since there is no currently available molecular marker that specifically delineates the OFT at these stages, we employed morphological landmarks to demarcate its boundaries (Fig. 1A,B; see Materials and Methods for more detail).

The enlargement of the OFT during its initial stages of assembly could be a cumulative outcome of changes in cell number, cell size and cell shape. To evaluate the degree to which changes in cell number accompany OFT assembly, we counted both endocardial and myocardial cells in the OFT at 36 and 51 hpf. Quantification of OFT endocardial cell number revealed that endocardial cells accrue as the OFT develops (Fig. 1C,G,S). Myocardial cell number in the OFT also increases during this time (Fig. 1K,O,W), consistent with previous studies examining the addition of late-differentiating cardiomyocytes to the arterial pole between 24 hpf and 48 hpf (de Pater et al., 2009; Lazic & Scott, 2011). Corresponding with the observed increases in the total number of cells, the number of endocardial and myocardial cells in a middle cross-section of the OFT also increases during this time interval (Fig. 1D,H,L,P,T,X).

To determine if changes in cell size and shape also contribute to OFT expansion between 36 and 51 hpf, we analyzed endocardial and myocardial cell surface area and circularity. We focused our analysis on the lateral walls of the OFT, which we refer to as the outer curvature (OC) and inner curvature (IC) due to their slightly convex and concave shapes at 51 hpf (Fig. 1A,B; see Materials and Methods for more detail). Our data demonstrate that OFT endocardial cells become smaller and rounder between 36 and 51 hpf (Fig. 1E,F,I,J,U,V). Simultaneously, and in contrast to the endocardial cells, myocardial cells in the OC become slightly larger, while also becoming less elongated (Fig. 1M,N,Q,R,Y,Z).

Taken together, our results indicate that OFT development between 36 and 51 hpf is accompanied by distinct patterns of cell behavior in both of its layers: in the endocardium, cell number increases over time and cells take on a more compact size and shape, while in the myocardium, cellular accrual is accompanied by regionalized expansion in surface area. How might these cellular behaviors underlie OFT growth? Given that endocardial cells are getting smaller and myocardial cells are enlarging only in a certain region, our data suggest cellular accrual as the primary mechanism for OFT expansion between 36 and 51 hpf. Importantly, our data indicate that the OFT widens during this interval, and that this increase in width corresponds with both myocardial and endocardial cellular accumulation along the circumference of the OFT.

### Proliferation and addition contribute to OFT endocardial expansion

Myocardial cell accrual in the OFT is thought to be a consequence of SHF differentiation (de Pater et al., 2009; Lazic & Scott, 2011), but little is known about how the OFT endocardium expands between 36 and 51 hpf. Since prior studies in zebrafish have indicated that ventricular endocardial cells proliferate between 24 and 48 hpf (Dietrich et al., 2014), we evaluated whether OFT endocardial cells proliferate between our stages of interest, using an EdU incorporation assay. At 51 hpf, we typically observed ∼5 EdU+ endocardial cells in the OFT, corresponding to an OFT endocardial proliferation index (PI) of 25.8 ± 1.2% (Fig. 2A-D). To complement this analysis, we performed a BrdU incorporation assay; these results revealed a similar PI in the OFT endocardium (28.9 ± 4.2 %; Fig. 2E-H). Together, our data suggest that proliferation is one mechanism by which endocardial cells accrue over time in the OFT.

**Figure 2.**
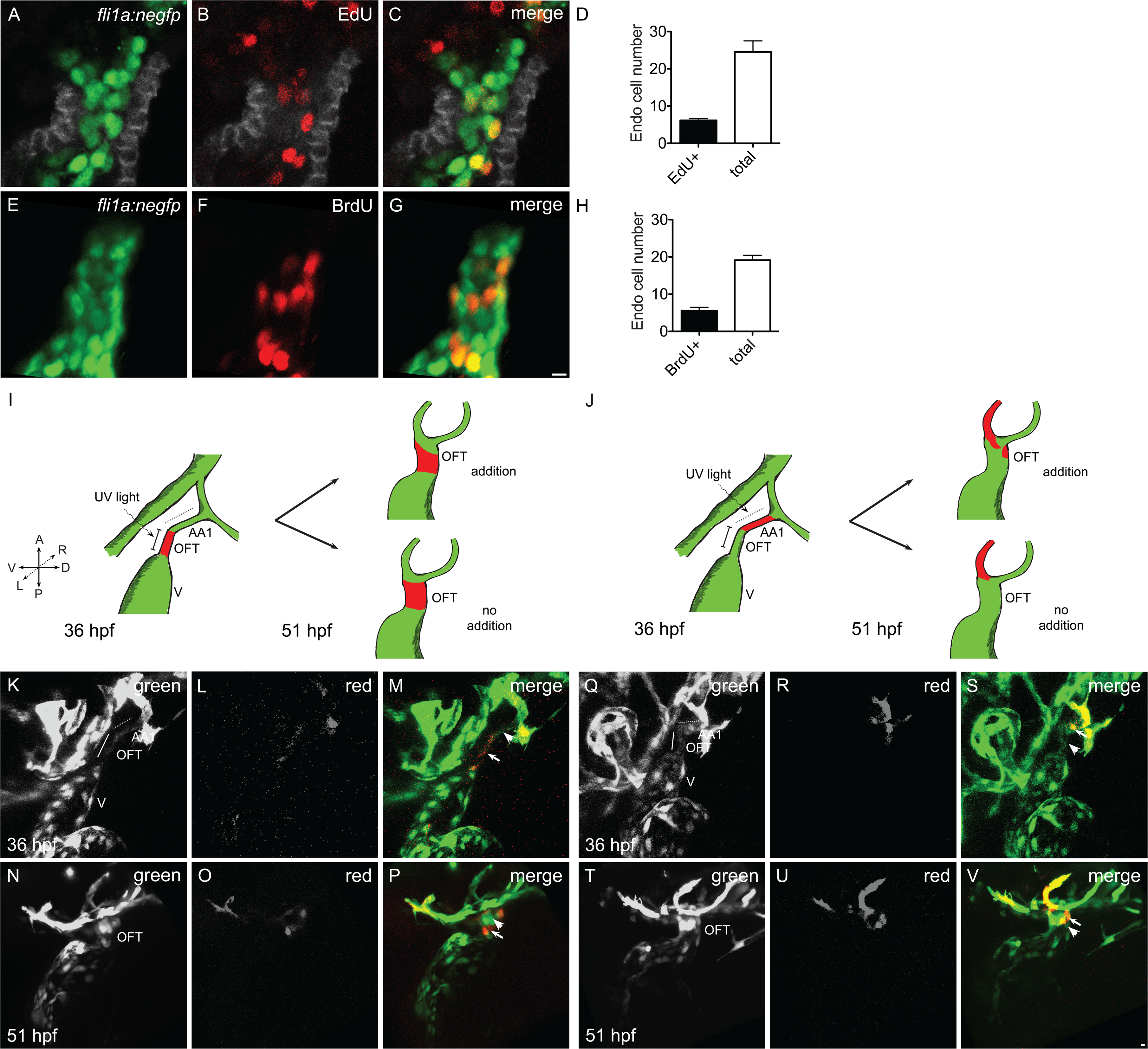
Endocardial proliferation and endothelial addition accompany OFT endocardial growth. **(A-D)** Lateral slices through the OFT at 51 hpf (A-C) show incorporation of EdU (red) into the *Tg(fli1a:negfp)*-labeled (green) endocardium between 36 and 51 hpf, with the myocardium marked by immunofluorescence for Alcama (grey). Quantification (D) indicates an OFT endocardial proliferation index (PI) of 25.8 ± 1.2% during this period of OFT expansion (n=6; N=1). **(E-H)** Lateral views of three-dimensional reconstructions at 51 hpf (E,F) show incorporation of BrdU (red) into the *Tg(fli1a:negfp)*-labeled (green) endocardium between 36 and 51 hpf. Quantification (H) indicates an OFT endocardial PI of 28.9 ± 4.2% (n=14; N=2). **(D,H)** Bar graphs indicate mean and SEM. **(I,J)** Cartoons illustrate the schemes for photoconversion experiments designed to assess the contribution of external sources (I) or endothelial cells from the first aortic arch (AA1) (J) to the OFT endocardium. **(K-P)** Three-dimensional reconstructions depict expression of *Tg*(*kdrl*:*dendra*), which drives the green-to-red photoconvertible protein Dendra in all endothelial cells. Photoconversion of the OFT endocardium (K-M) at 36 hpf was followed by detection of green cells, which had not been labeled by photoconversion, in the OFT endocardium at 51 hpf (N-P), indicating the contribution of cells from external sources to the OFT endocardium during this time period (n=3; N=2). Arrowheads indicate examples of cells that were not labeled by photoconversion (green only), and arrows indicate examples of cells that were labeled by photoconversion (red and green). **(Q-V)** Photoconversion of a portion of AA1 endothelium (red) at 36 hpf (Q-S) was followed by detection of labeled cells in the OFT at 51 hpf (T-V) (n=5; N=2). Scale bars: 5 µm.

Prior studies in zebrafish have also determined that cells from the arterial vasculature can migrate into the heart, in a direction opposite to that of blood flow, between 24 and 48 hpf (Rochon et al., 2016). We therefore investigated whether endothelial cells from external sources supply cells to the OFT endocardium between 36 and 51 hpf. In these experiments, we employed embryos carrying the transgene *Tg*(*kdrl*:*dendra*), in which the green-to-red photoconvertible protein Dendra is expressed in the endothelium. First, we employed photoconversion to label cells throughout the OFT endocardium at 36 hpf (Fig. 2I,K-M), followed by subsequent evaluation of the composition of the OFT. By 51 hpf, we found that the OFT contained two populations of cells: those with only the native green form of Dendra, and those with the photoconverted red form of Dendra in addition to the newly synthesized green form (Fig. 2I,N-O). The appearance of “green-only” cells in the OFT suggests that the addition of externally derived cells contributes to the growth of the OFT endocardium. Considering the proximity of the first aortic arch (AA1) to the OFT, we next tested whether cells from AA1 come to occupy the OFT. Specifically, we labeled a portion of AA1 via photoconversion at 36 hpf (Fig. 2J,Q-S) and then evaluated the subsequent locations of the labeled cells. Indeed, we found labeled cells in the OFT endocardium by 51 hpf (Fig. 2J,Q-S), most frequently in the distal OFT (Fig. S2-1), suggesting that cells from AA1 relocate to join the OFT endocardium. Thus, our experiments show that both endocardial proliferation and addition of AA1 cells contribute to the accrual of cells in the OFT endocardium.

The observed patterns of cellular accrual within the OFT correspond to the changes in OFT tissue morphology that take place between 36 and 51 hpf. Three-dimensional surface reconstructions of the OFT endocardial walls (see Materials and Methods) demonstrated that, at 36 hpf, the OFT endocardium is a relatively flattened tube that appears uniformly wide along its proximal-distal axis (Fig. 3A-E,X; Fig. S3-1; Video S1). However, by 51 hpf, the OFT adopts a funnel-like shape, such that its distal opening into the aortic arches is larger than its proximal connection to the ventricle (Fig. 3K-O,Y; Fig. S3-1; Video S2). In addition, the enclosed volume of the OFT endocardium expands substantially between 36 and 51 hpf (Fig. 3U); this expansion was evident both in fixed samples and in live embryos (Figs. 3U and S3-2). Additionally, the enclosed volume of the OFT myocardium increases between 36 and 51 hpf (Fig. 3V); the relatively consistent ratio of myocardial and endocardial volumes at 36 and 51 hpf (Fig. 3W) suggests coordinated morphogenesis of the two layers. Thus, endocardial cellular accrual coincides with an expansion of both endocardial and myocardial volume, which presumably facilitates OFT lumen capacity as blood flow increases over time.

**Figure 3.**
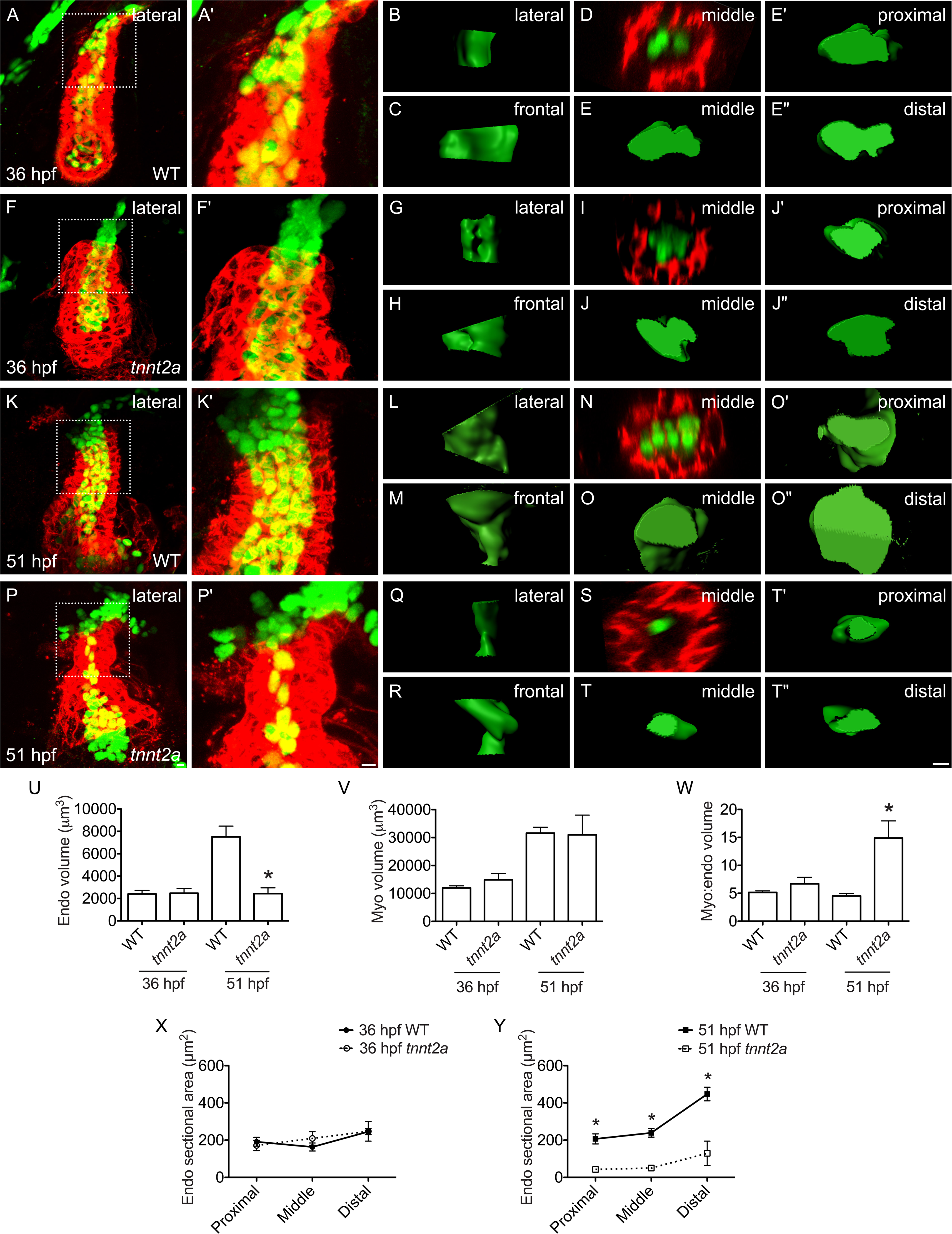
Cardiac function promotes OFT growth. **(A,F,K,P)** Lateral views of three-dimensional reconstructions compare the OFT and ventricle in wild-type (WT) (A,K) and *tnnt2a* mutant embryos (F,P), labeled by immunofluorescence for Alcama (red) and *Tg(fli1a:negfp)* (green). Since Alcama is found in the basolateral boundaries of myocardial cells, the endocardium is visible through the myocardial layer. Closer views of the OFT (A’,F’,K’,P’) demonstrate relatively similar morphology in WT (A’) and *tnnt2a* mutant (F’) embryos at 36 hpf (n=5,4; N=2). See also Videos S1 and S3. However, by 51 hpf, whereas the WT OFT endocardial and myocardial volumes (K’) have expanded significantly (p<0.0005, Student’s T test), the OFT endocardium in *tnnt2a* mutants (P’) has failed to expand, as quantified in (U,V), resulting in a significantly higher ratio of myocardial to endocardial volume in *tnnt2a* mutants by 51 hpf (W) (n=7,6; N=2; asterisks depict significant differences from wild-type at same stage, p<0.005, Student’s T test). See also Videos S2 and S4. **(B,C,G,H,L,M,Q,R)** Lateral (B,G,L,Q) and frontal (C,H,M,R) views of smoothed OFT endocardial surfaces generated from three-dimensional reconstructions of cytoplasmic GFP localization in *Tg*(*kdrl*:*grcfp*) embryos (Fig. S3-1) display the relatively similar OFT morphologies in WT (B,C) and *tnnt2a* mutants (G,H) at 36 hpf and the different OFT morphologies in WT (L,M) and *tnnt2a* mutants (Q,R) at 51 hpf. Lateral and frontal surfaces are oriented with their proximal end at the bottom. Of note, the lateral side of the WT OFT at 36 hpf (B) appears to transition into the frontal side at 51 hpf (M), suggesting that the OFT rotates 90° during this time. **(D,E,I,J,N,O,S,T)** Middle cross-sections through the OFTs shown in (A’,F’,K’,P’) indicate that the OFT endocardium collapses in *tnnt2a* mutants between 36 (I) and 51 hpf (S), in contrast to its broadening in WT (D,N). These contrasts are also evident in comparisons between middle (E,J,O,T), proximal (E’,J’,O’,T’) and distal (E’’,J’’,O’’,T’’) cross-sections through the smoothed surfaces of the OFT endocardium in *Tg*(*kdrl*:*grcfp*)-expressing embryos (Fig. S3-1), as quantified in (X,Y) (n=6, 4 for 36 hpf; 5,5 for 51 hpf; N=2; asterisks depict significant differences from WT at same positions, p<0.005, Student’s T test). The broadening of the WT endocardium between 36 and 51 hpf is especially striking at its distal end (E’’,O’’,X,Y) (p<0.05, Student’s T test). Proximal, middle, and distal sections are oriented with the OC of the OFT to the left. **(U-Y)** Bar graphs and line graphs indicate mean and SEM. Scale bars: 5 µm.

### Cardiac function promotes OFT endocardial expansion

Since the OFT is assembled atop a beating heart, we hypothesized that functional cues could influence the morphogenesis of the OFT endocardium. To begin testing this, we analyzed OFT endocardial dimensions in *troponin T type 2a* (*tnnt2a*) mutant embryos (also known as *silent heart* mutants), which lack cardiac contractility and, consequently, blood flow (Sehnert et al., 2002). At 36 hpf, we found that OFT endocardial volume and cross-sectional areas are indistinguishable between wild-type and *tnnt2a* mutant embryos (Fig. 3A-J,U,X; Videos S1 and S3). However, by 51 hpf, while the wild-type OFT endocardium grows in volume and distal sectional area, the *tnnt2a* mutant OFT endocardium fails to expand (Fig. 3K-T,U,X,Y; Videos S2 and S4). Similar to our observations in fixed samples, our analysis of live embryos highlighted that the OFT endocardium in *tnnt2a* mutants appears as a collapsed, thin column of cells that encloses a much smaller volume than the wild-type endocardium (Fig. S3-2). In contrast, the OFT myocardial volume is relatively similar in wild-type and *tnnt2a* mutant embryos (Fig. 3V). Together, these results demonstrate that cardiac function plays an essential role in promoting the expansion of the OFT endocardium. In this way, cardiac function appears to feed back to the OFT, influencing its morphogenesis and, consequently, cardiac output.

We next investigated the cellular mechanisms through which cardiac function regulates OFT endocardial expansion. First, we examined OFT endocardial cell number in *tnnt2a* mutants. At 36 hpf, the number of OFT endocardial cells in wild-type and *tnnt2a* mutant embryos is comparable (Fig. 4A,C,K,L), consistent with the similarity in OFT endocardial volume at this stage. By 51 hpf, however, the number of OFT endocardial cells is significantly smaller in *tnnt2a* mutants than in wild-type (Fig. 4B,D,K,L). Additionally, although *tnnt2a* mutants have a normal number of OFT myocardial cells at 51 hpf, their number of cardiomyocytes in a middle cross-section is reduced (Fig S4-1). Finally, in addition to the failure of OFT endocardial accumulation in *tnnt2a* mutants, we found that the OFT endocardial cells in *tnnt2a* mutants stay relatively large and elongated at 51 hpf, rather than becoming smaller and more circular as in wild-type (Fig. S4-2). Altogether, our data suggest that failure to accumulate an appropriate number of endocardial cells underlies the observed collapse of the OFT endocardium in *tnnt2a* mutants.

**Figure 4.**
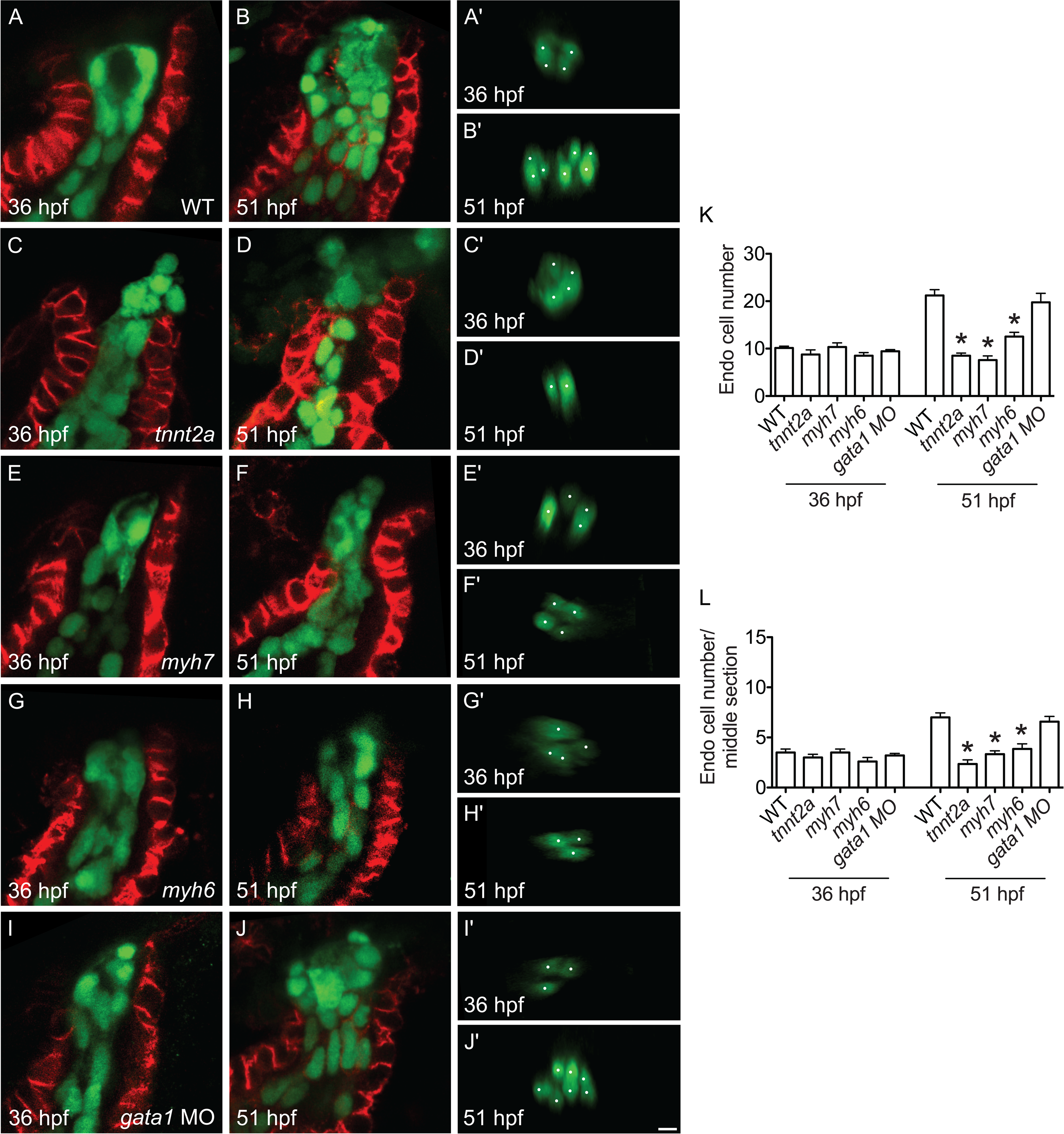
Cardiac function promotes accrual of endocardial cells in the OFT. **(A-J)** Lateral slices (A-J) and middle cross-sections (A′-J′) through the OFT (as in Fig. 1C,D) illustrate the number of OFT endocardial cells at 36 (A,C,E,G,I) and 51 hpf (B,D,F,H,J). As shown in Fig. 1S,T, the number of endocardial cells in the WT OFT increases between 36 (A,A’) and 51 hpf (B,B’), as quantified in (K,L) (p<0.0001, Student’s T test). In contrast, in *tnnt2a* mutant embryos, OFT endocardial cells fail to accrue between 36 (C,C′) and 51 hpf (D,D′). A similar deficiency in OFT endocardial accrual is found in *myh7* mutant embryos (E,E′,F,F′), and *myh6* mutant embryos (G,G′,H,H′) also exhibit a reduction in OFT endocardial cell number by 51 hpf. However, *gata1* morphants (I,I′,J,J′) exhibit a normal number of OFT endocardial cells. **(K,L)** Bar graphs indicate mean and SEM (n=15, 8, 6, 6, 7 for 36 hpf; 24, 6, 10, 13, 8 for 51 hpf; N=2; asterisks represent significant differences from WT at same stage, p<0.05, Student’s T test.) Scale bar: 5 µm.

To evaluate whether the *tnnt2a* mutant phenotype reflects the consequences of a lack of cardiac function, rather than a lack of Tnnt2a specifically, we next examined the OFT endocardium in *myh7* mutants, which lack ventricular contractility and exhibit substantially reduced blood flow, due to a lack of the ventricular myosin heavy chain Myh7 (also known as Vmhc) (Auman et al., 2007). Strikingly, the phenotype of the OFT endocardium in *myh7* mutants resembles the phenotype observed in *tnnt2a* mutants, with a significant failure to increase OFT endocardial cell number between 36 and 51 hpf in each scenario (Fig. 4C-F,K,L). Taken together, these data suggest that cardiac function is crucial for promoting the accrual of OFT endocardial cells.

Our analyses of *tnnt2a* and *myh7* mutants do not allow us to distinguish between the potential effects of contractility and blood flow on OFT expansion. To evaluate the effects of altering blood flow without interfering with the contractility of the ventricle and the OFT, we assessed the phenotype of *myh6* mutants, which lack the atrial myosin heavy chain Myh6 (also known as Amhc) (Berdougo et al., 2003). In *myh6* mutants, the ventricle contracts at a normal rate while the atrium is non-contractile, and blood flow through the heart is relatively inefficient, as indicated by a characteristic pooling of blood near the inflow tract (Berdougo et al., 2003). As in *tnnt2a* and *myh7* mutants, OFT endocardial cell number is normal in *myh6* mutants at 36 hpf (Fig. 4G,K,L). However, at 51 hpf, the *myh6* mutant OFT contains significantly fewer endocardial cells in comparison to wild-type (Fig. 4H,K,L). Thus, loss of atrial function impedes the accrual of OFT endocardial cells, implying that patterns of blood flow influence OFT morphogenesis. However, circulating erythrocytes seem irrelevant to OFT endocardial growth, since *gata1* morpholino-injected embryos (“morphants”) lack erythrocytes (Vermot et al., 2009) and exhibit normal numbers of OFT endocardial cells (Fig. 4I-L). Thus, our data suggest the possibility that particular flow parameters, separate from the shear forces generated by erythrocytes, act independently of contractile stretch to promote OFT endocardial expansion.

### Cardiac function regulates both endocardial proliferation and endothelial addition to the OFT

We next examined whether functional cues regulate either endocardial proliferation or endothelial addition to the OFT. First, we inspected OFT endocardial proliferation in *tnnt2a* mutants using both EdU and BrdU incorporation assays (Figs. 5 and S5-1). Strikingly, whereas endocardial cells in the wild-type OFT exhibit a PI of 29.6 ± 3.6% between 36 and 51 hpf, the PI in the *tnnt2a* mutant OFT endocardium is significantly reduced to 10.2 ± 4.3% (Figs. 5A-F,M and S5-1A-D,I). This reduction is recapitulated in the *myh7* mutant OFT endocardium, which exhibits a PI of 15.6 ± 4.8% (Figs. 5G-I,M and S5-1E,F,I). Thus, cardiac function drives proliferation of the OFT endocardium, as has been previously suggested for the ventricular endocardium (Dietrich et al., 2014).

**Figure 5.**
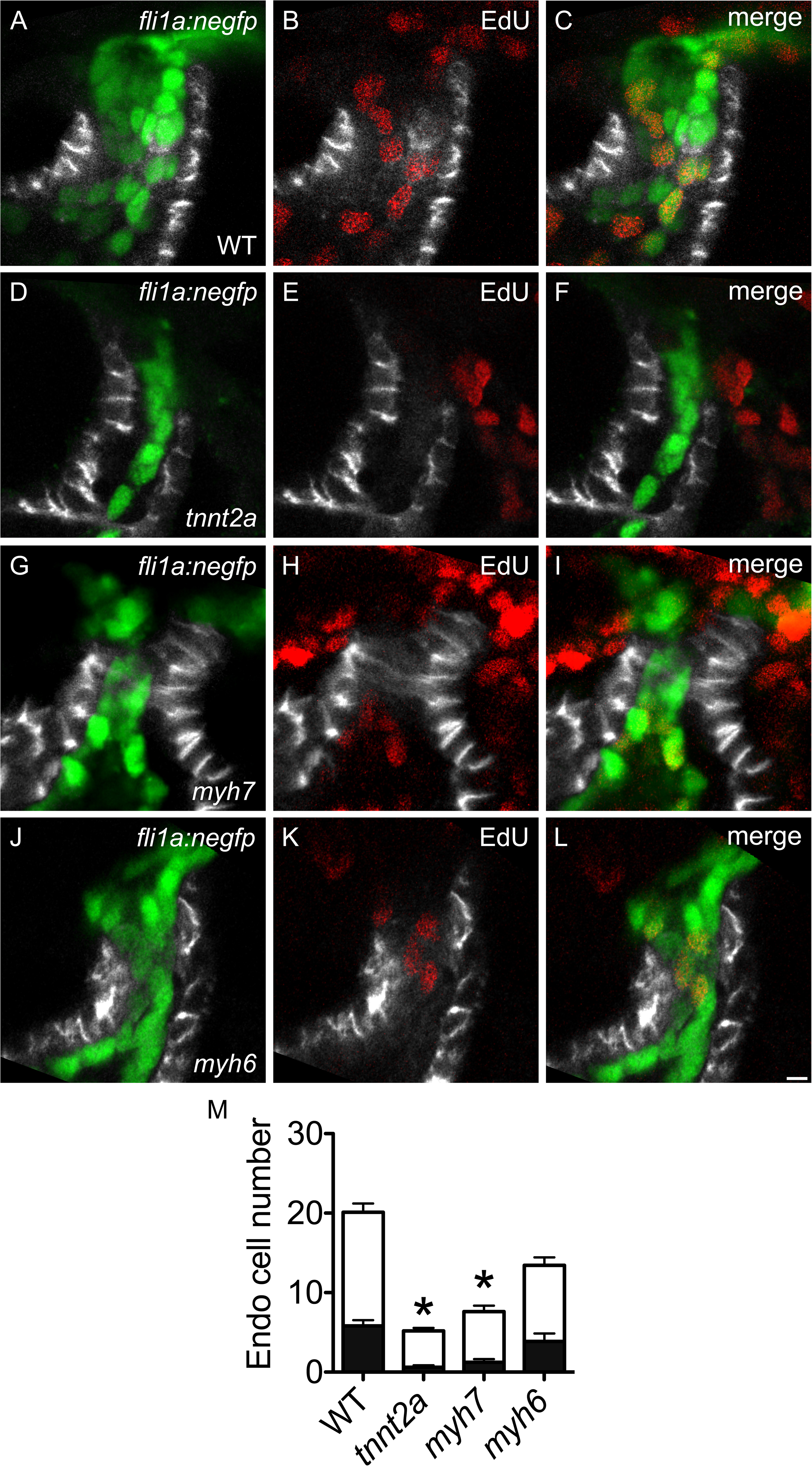
OFT endocardial proliferation depends upon functional inputs. **(A-L)** Lateral slices through the OFT at 51 hpf (as in Fig. 2A-C) illustrate EdU incorporation between 36 and 51 hpf in WT (A-C), *tnnt2a* mutant (D-F), *myh7* mutant (G-I) and *myh6* mutant (J-L) embryos. **(M)** Black and white bars in graph represent mean number of EdU+ and EdU- endocardial cells, respectively, with SEM. EdU incorporation is significantly reduced in *tnnt2a* mutants (PI=10.2 ± 4.3%) and *myh7* mutants (PI=15.6 ± 4.9%), but not in *myh6* mutants (PI=27.8 ± 6.3%), compared to WT (PI=29.6 ± 3.7%) (n=17, 10, 13, 7; N=2; asterisks indicate significant differences in PI compared to WT, p<0.05, Student’s T test). Scale bar: 5 µm.

Next, since prior studies had shown that cardiac function can promote migration of endothelial cells toward the heart between 24 and 48 hpf (Rochon et al., 2016), we evaluated whether cardiac function also influences endothelial displacement from AA1 into the OFT between 36 and 51 hpf, using photoconversion to label AA1 cells in *tnnt2a* mutant embryos carrying *Tg*(*kdrl*:*dendra*). In contrast to our observations in wild-type siblings, in which labeled AA1 cells consistently contributed to the OFT endocardium, we rarely found labeled cells in the OFT endocardium of *tnnt2a* mutants (Figs. 6A-L,S,T; Videos S5 and S6). Similar observations were obtained in *tnnt2a* morphants (Fig. S6-1).

**Figure 6.**
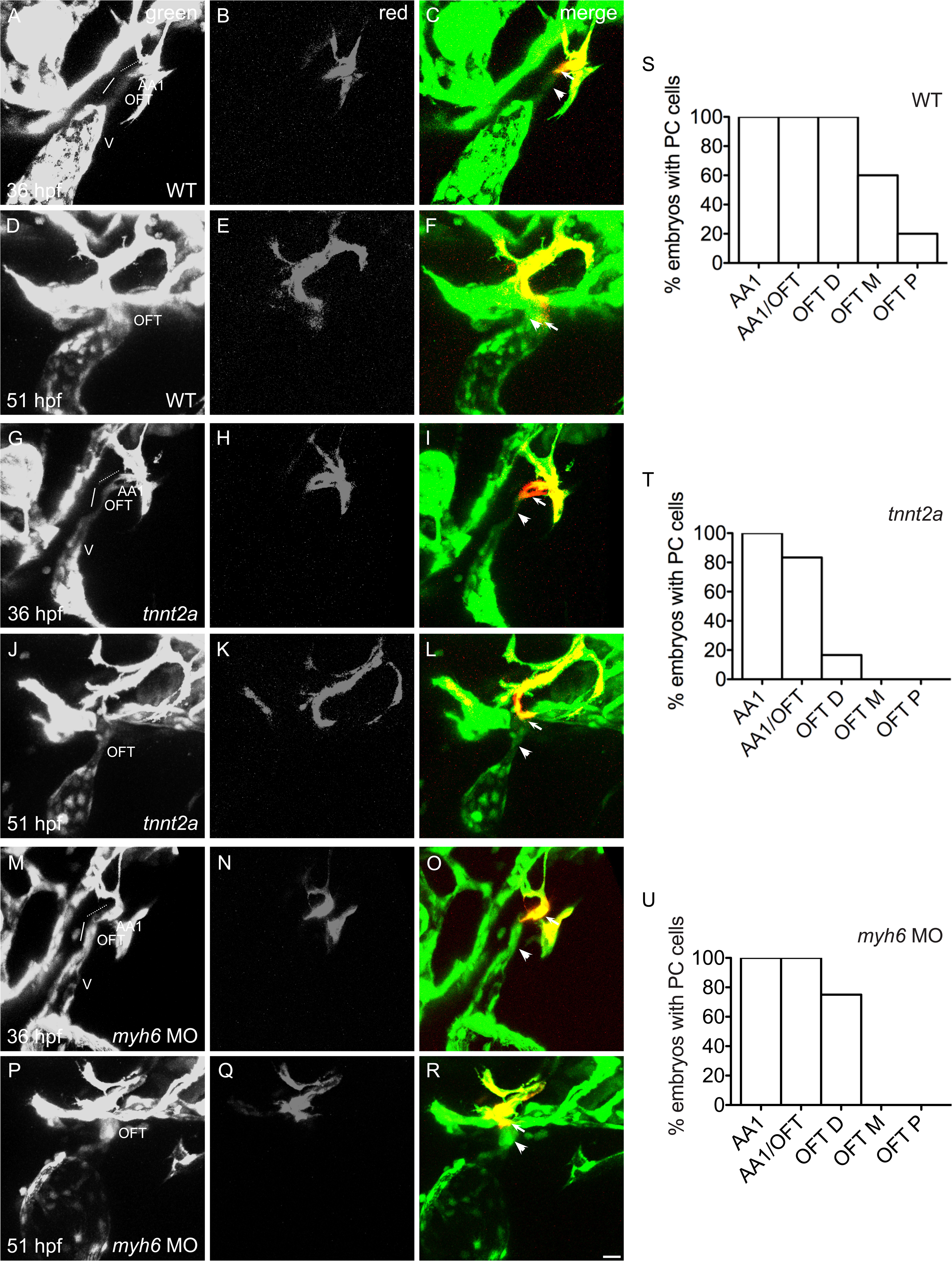
Cardiac function promotes addition of endothelial cells from the aortic arches to the OFT endocardium. **(A-R)** Three-dimensional reconstructions depict the photoconversion of a portion of AA1 endothelium at 36 hpf (as in Fig. 2Q-S) in WT (A-C), *tnnt2a* mutants (G-I) or *myh6* morphants (M-O). By 51 hpf, labeled cells are routinely detected (as in Fig. 2T-V) in the WT OFT, as quantified in (S) (n= 5; N=2). See also Video S5. In contrast, in *tnnt2a* mutants (J-L) and *myh6* morphants (P-R) at 51 hpf, labeled cells are found less often in the distal OFT and are entirely absent from the middle and proximal OFT, as quantified in (T,U) (n=6 for *tnnt2a*, 5 for *myh6* MO; N=2). See also Videos S6 and S7. Arrowheads indicate examples of cells that were not labeled by photoconversion (green only), and arrows indicate examples of cells that were labeled by photoconversion (red and green). **(S-U)** Bar graphs (as in Fig. S2-1) indicate the percentage of embryos in which labeling of AA1 endothelium at 36 hpf resulted in detection of photoconverted (PC) cells in AA1, AA1/OFT, OFT D, OFT M, or OFT P at 51 hpf. Scale bar: 20 µm.

To determine if either or both of these cellular processes depend on atrial function, we inspected endocardial proliferation and endothelial addition in the *myh6* mutant OFT. We were intrigued to find that OFT endocardial cells in *myh6* mutants exhibit a normal PI of 27.8 ± 6.3% (Figs. 5J-M and S5-1G-I). In contrast, endothelial addition to the OFT is reduced in *myh6* morphants when compared to wild-type (Fig. 6M-R,U; Video S7). Altogether, whereas the *tnnt2a* and *myh7* mutant phenotypes indicate that both endocardial proliferation and endothelial addition are dependent on cardiac function, the effects of *myh6* loss-of-function suggest that endothelial addition, but not necessarily endocardial proliferation, requires cues generated by blood flow.

### Klf2a and Klf2b are not essential for endocardial expansion in the OFT

Downstream of the biomechanical cues generated by cardiac function, what are the mechanosensitive pathways that instigate changes in OFT cell behavior? We first considered whether Klf2 transcription factors might regulate cellular accrual in the OFT. *klf2a* is expressed in the endocardium at 36 hpf, and its expression becomes markedly upregulated in the OFT endocardium by 48 hpf (Vermot et al., 2009). Prior studies in both cultured cells (Dekker et al., 2002) and murine embryos (Goddard et al., 2017) have suggested that endothelial *Klf2* expression is dependent upon shear forces. In zebrafish, *klf2a* expression in the endocardium is also responsive to reversing flows (Vermot et al., 2009); specifically, endocardial *klf2a* expression is diminished in *tnnt2a* mutants, which lack cardiac function, and *gata2* morphants, which exhibit reduced retrograde flow (Vermot et al., 2009). Additionally, *klf2a* appears to promote endocardial cell accumulation in the atrioventricular canal (Vermot et al., 2009) and to drive restriction of endocardial cell size during ventricular chamber expansion (Dietrich et al., 2014).

To examine the roles of *klf2a* and its paralog *klf2b* in OFT assembly, we examined OFT endocardial cell number in embryos homozygous for mutations in both *klf2a* and *klf2b* (henceforth called *klf2* mutants) (Rasouli et al., 2018) at 51 hpf. OFT endocardial cell number in ten out of eleven *klf2* mutants appeared remarkably similar to their control siblings (Fig.S7-1A,B,D), suggesting that Klf2 factors do not govern OFT endocardial expansion. We did, however, observe collapsed endocardial morphology and a significant reduction in OFT endocardial cell number in one out of eleven *klf2* mutant embryos (Fig. S7-1C), suggesting the possibility of a poorly penetrant phenotype; similarly, defects in atrioventricular canal morphogenesis also appeared with ∼10% penetrance in *klf2* mutants by 48 hpf (Rasouli et al., 2018). Thus, although we cannot rule out some contribution of Klf2 factors to OFT endocardial expansion, it seems that they are not essential.

### Acvrl1 promotes OFT expansion via endothelial addition, but is not required for endocardial proliferation

We next evaluated another interesting candidate for a mechanosensitive mediator of OFT endocardial development, the TGFβ receptor Acvrl1. Expression of *acvrl1* in the arterial endothelium, particularly in AA1, is coincident with the initiation of blood flow around 26 hpf, and perdures until at least 48 hpf (Corti et al., 2011). Importantly, *acvrl1* expression is significantly reduced in *tnnt2a* mutants, suggesting a potent influence of cardiac function on Acvrl1-mediated endothelial cell behaviors (Corti et al., 2011). Indeed, like *tnnt2a* morphants, *acvrl1* mutants demonstrate reduced endothelial migration toward the heart between 24 hpf and 48 hpf (Rochon et al., 2016).

To examine the influence of Acvrl1 on OFT endocardial morphogenesis, we first evaluated OFT endocardial cell number in *acvrl1* morphants (Corti et al., 2011). Previous studies have established that *acvrl1* morphants phenocopy *acvrl1* mutants; for example, both mutant and morphant embryos exhibit characteristic enlargement of cranial vessels by 36 hpf (Corti et al., 2011; Laux et al., 2013; Rochon et al., 2016). At 36 hpf, we found a comparable number of OFT endocardial cells in uninjected controls and *acvrl1* morphants (Fig. 7A,C,D). However, OFT endocardial cells do not accumulate normally in *acvrl1* morphants, resulting in a significant reduction of OFT endocardial cell number by 51 hpf (Fig. 7B,C,E), and *acvrl1* mutants exhibit a similar defect in the number of OFT endocardial cells (Fig. S7-2). Notably, the failure of endocardial accrual in *acvrl1* morphants and mutants (Figs. 7C and S7-2C) resembles that observed in *tnnt2a*, *myh7*, and *myh6* mutants (Fig. 4K); yet, in contrast to the contractility and blood flow defects in these mutants, *acvrl1* morphants display relatively normal cardiac function between 36 and 51 hpf, with the exception of a slight decline in heart rate by 51 hpf (Fig. S7-3). Thus, *acvrl1* is required to mediate OFT endocardial expansion, even in the context of relatively normal cardiac function.

**Figure 7.**
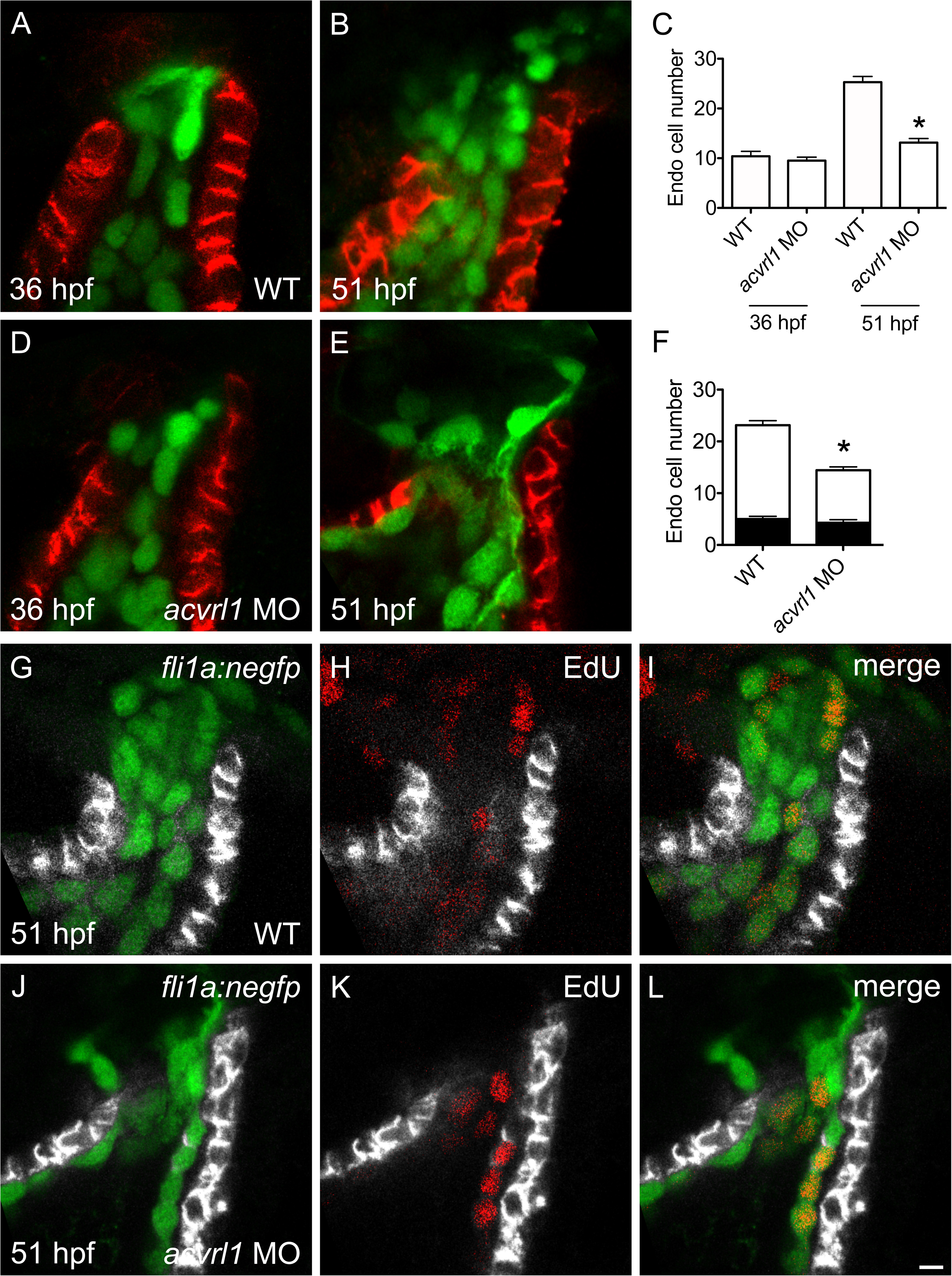
Acvrl1 promotes OFT endocardial expansion, but is not required for endocardial proliferation. **(A-E)** Lateral slices (A,B,D,E) through the OFT (as in Fig. 1C) illustrate the number of OFT endocardial cells at 36 (A,D) and 51 (B,E) hpf. At 36 hpf, WT control (A) and *acvrl1* morphant (D) embryos have similar numbers of OFT endocardial cells (n=5, 8; N=2). However, by 51 hpf, *acvrl1* morphants (E) have significantly fewer OFT endocardial cells compared to WT controls (B) (n=14, 16; N=2). Bar graph (C) indicates mean and SEM (asterisk indicates significant difference from WT controls, p<0.0001, Student’s T test). **(F-L)** Lateral slices (G-L) through the OFT at 51 hpf (as in Fig. 2A-C) illustrate EdU incorporation between 36 and 51 hpf in WT control (G-I) and *acvrl1* morphant (J-L) embryos. Black and white bars in graph (F) represent mean number of EdU+ and EdU- endocardial cells, respectively, with SEM. The reduced number of OFT endocardial cells in *acvrl1* morphants is not a consequence of defective proliferation, since the number of EdU+ cells in the *acvrl1* morphant OFT (J-L) is similar to that in WT controls (G-I), resulting in an *acvrl1* morphant PI of 29.1 ± 2.7%, which is significantly higher than that of WT controls (PI=21.5 ± 1.9 %) (n= 8, 7; N=2; asterisk indicates significant difference in PI from WT controls, p<0.05, Student’s T test). Scale bar: 5 µm.

To elucidate the cellular mechanisms that underlie the reduction in endocardial cell number in *acvrl1* morphants, we assayed both the proliferation of OFT endocardial cells and the addition of endothelial cells from AA1 to the OFT. First, using an EdU incorporation assay, we found that *acvrl1* morphants exhibit a PI of 29.1 ± 2.7%, in comparison to 21.5 ± 1.9% in uninjected control embryos (Fig. 7F-L), indicating that Acvrl1 is not required for OFT endocardial proliferation. In contrast, endothelial addition from AA1 into the OFT was substantially inhibited in *acvrl1* morphants (Fig. 8, Video S8). This observation is consistent with the known role of Acvrl1 in promoting endothelial migration (Rochon et al., 2016), and, importantly, it reveals a previously unappreciated function for Acvrl1 in regulating endocardial morphogenesis. Synthesizing these data with the reduced *acvrl1* expression seen in *tnnt2a* mutants (Corti et al., 2011) and our observations of decreased endothelial addition to the OFT in *tnnt2a* and *myh6* mutants (Fig. 6T,U), Acvrl1 emerges as an important mediator of OFT expansion through its regulation of endothelial cell displacement into the OFT, potentially in response to cues generated by blood flow.

**Figure 8.**
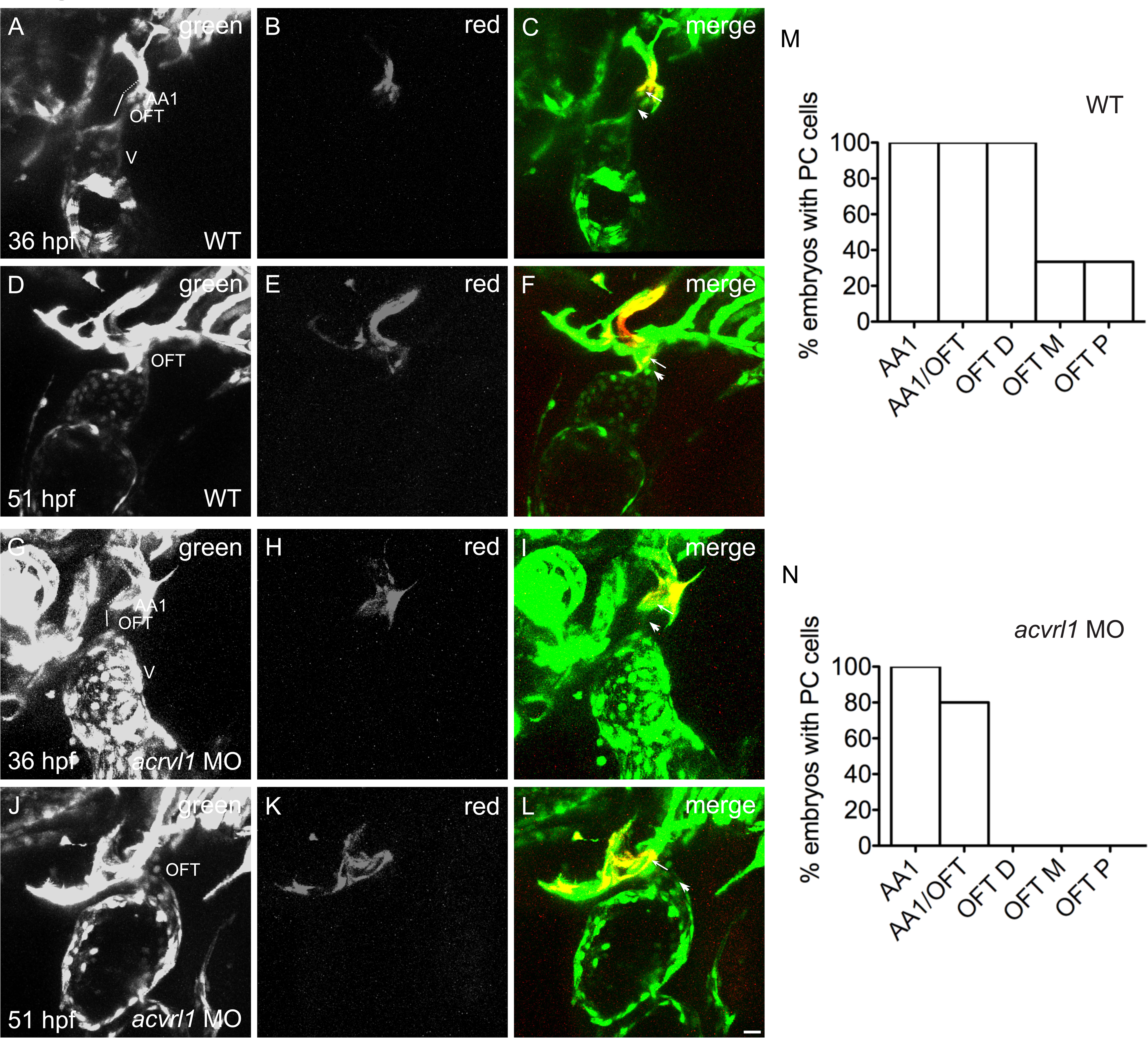
Acvrl1 promotes OFT growth by regulating endothelial addition. **(A-L)** Three-dimensional reconstructions depict the photoconversion of a portion of AA1 endothelium at 36 hpf (as in Fig. 2Q-S) in WT embryos (A-C) or *acvrl1* morphants (G-I). By 51 hpf, labeled cells are routinely detected (as in Fig. 2T-V) in the WT OFT (D-F), as quantified in (M) (n= 3; N=2). However, in *acvrl1* morphants, labeled AA1 endothelial cells fail to accrete to the OFT by 51 hpf (J-L) (n=5; N=2). See also Video S8. Arrowheads indicate examples of cells that were not labeled by photoconversion (green only), and arrows indicate examples of cells that were labeled by photoconversion (red and green). **(M,N)** Bar graphs (as in Fig. S2-1) indicate the percentage of embryos in which photoconversion of AA1 endothelium at 36 hpf resulted in detection of labeled cells in AA1, AA1/OFT, OFT D, OFT M, or OFT P at 51 hpf. Scale bar: 20 µm.

## DISCUSSION

Our studies provide new perspectives on the essential mechanisms through which the OFT attains its stereotypical morphology. By capturing the cellular dynamics underlying OFT assembly, we have found that the expansion of the OFT endocardium involves both the proliferation of endocardial cells and the incorporation of endothelial cells from AA1 into the OFT. Additionally, we have demonstrated that cardiac function plays a crucial role in promoting OFT endocardial expansion via both proliferation and cell addition. Finally, we have uncovered a previously unappreciated function of Acvrl1 in supporting OFT endocardial growth by driving displacement of endothelial cells from AA1 into the OFT. Altogether, these findings suggest a novel model in which biomechanical cues generated by cardiac function promote both endocardial proliferation and Acvrl1-dependent endothelial addition in order to forge the specific endocardial dimensions of the OFT lumen, thereby creating an appropriate conduit for circulation and serving as a platform for myocardial organization.

Our data suggest that proliferation and cell addition collaboratively shape the OFT endocardium, but they do not definitively delineate whether these two cell behaviors are independent or coupled. Our observations of the *myh6* and *acvrl1* loss-of-function phenotypes, in which endothelial cell addition is significantly reduced while the endocardial proliferation index is not, suggest that endocardial proliferation and endothelial cell addition are separable processes that are regulated independently. Since the endothelial cells that are displaced from AA1 are routinely found in the distal portion of the OFT, it is appealing to envision that proliferation and cell addition could involve separate sets of cells: a relatively proximal population that is proliferative and sets the proximal diameter of the OFT and a relatively distal population that enters from AA1 and facilitates the distal expansion of the OFT. Even in this scenario, however, there could be interdependence of the two populations; for example, addition of endothelial cells from AA1 could potentially depend upon the prerequisite presence of proliferating endocardial cells in the proximal OFT. Precise articulation of the relationship between endocardial proliferation and endothelial addition into the OFT will require high-resolution time-lapse analysis of endocardial cell division and movement, as well as the identification of mutants or tools that could selectively interfere with endocardial proliferation in a spatially and temporally controlled fashion.

If endocardial proliferation and endothelial cell addition are indeed uncoupled, they may also be governed by distinct biomechanical mechanisms. For example, fluid parameters, including shear forces, blood pressure or oscillatory shear indices, might influence cellular behaviors independent of the tensile forces associated with contractility. Our data indicate that cell addition, but not proliferation, is hampered when atrial function is disrupted by *myh6* loss-of-function; in this scenario, patterns of blood flow are aberrant, while ventricular and OFT contractility are preserved. This is in contrast to our observations in *tnnt2a* mutants, in which both blood flow and OFT contractility are impaired, and both cell addition and proliferation are inhibited. These results suggest the possibility that fluid forces act to stimulate endothelial addition to the OFT, while tensile forces promote endocardial proliferation. Alternatively, fluid forces could regulate both cell addition and proliferation: perhaps the relatively modest flow defects in *myh6* mutants are inadequate to interfere with the proliferation of OFT endocardial cells.

In evaluating the specific fluid forces that could influence OFT endocardial expansion, we note that the normal OFT phenotype observed in *gata1* morphants suggests that shear stress generated by erythrocytes is not a key factor. This is in agreement with previous studies in the endothelium (Corti et al., 2011), atrioventricular canal (Heckel et al., 2015; Vermot et al., 2009) and ventricle (Dietrich et al., 2014), where loss of *gata1* was not found to influence flow-induced morphogenetic mechanisms. What other flow-related parameters could instigate OFT endocardial growth? Perhaps the shear forces generated by blood plasma are sufficient to trigger changes in endocardial cell behavior. Flow direction may also be a key influence on endocardial cell accrual; in the atrioventricular canal, reversing flows are thought to stimulate endocardial cell accumulation in the atrioventricular canal (Vermot et al., 2009). Another intriguing possibility is that particular patterns of blood pressure could stretch endocardial cells in a manner that provokes specific cellular responses. In future work, a combination of techniques that directly manipulate flow direction or induce stretch (Anton et al., 2013; Marjoram et al., 2016; Sidhwani & Yelon, 2019), together with controlled flow environments to house explanted hearts (Cao & Poss, 2016; Wong et al., 2016), could enable the dissection of potential influences of specific functional cues on OFT morphogenesis.

In addition to considering flow-induced forces as regulators of endocardial expansion, it is important to consider blood as a carrier of vital molecules. For example, blood flow may transport ligands to activate the signaling pathways that promote endothelial cell addition to the OFT. Indeed, the Acvrl1 ligand Bmp10 is known to be in circulation (Laux et al., 2013; Ricard et al., 2012) and is thought to be transported by blood flow from the heart to the site of *acvrl1* expression in the arterial vessels (Laux et al., 2013). Thus, we can envision a scenario in which blood flow regulates both *acvrl1* expression as well as *bmp10* dissemination (Corti et al., 2011; Laux et al., 2013; Rochon et al., 2016) to facilitate displacement of endothelial cells from AA1 into the OFT.

This role of the Acvrl1 pathway has the potential to be conserved in mammals: it is intriguing to note that mice lacking the Acvrl1 coreceptor Endoglin exhibit misshapen OFT morphology (Arthur et al., 2000). The function of Acvrl1 during OFT morphogenesis has not been examined directly in mice, and it is not known whether the instances of cardiac failure in mouse *Acvrl1* mutants (Morine et al., 2017a; Morine et al., 2017b) or in human HHT2 patients (Cho et al., 2012; Goussous et al., 2009) are associated with defective cardiac development. Since our studies assert that Acvrl1 is a key mediator of endothelial displacement during OFT endocardial expansion in zebrafish, it will be worthwhile for future studies to investigate whether any early defects in OFT development appear in mice lacking Acvrl1.

While our work provides new insights into the molecular regulation of endothelial addition to the OFT, we do not yet know the molecular pathways by which cardiac function influences OFT endocardial proliferation. Previous studies have shown that endocardial proliferation in the ventricle similarly relies on functional cues (Dietrich et al., 2014), but the mechanosensitive regulators remain unidentified in this context as well. Recent work suggest that endocardial proliferation in the atrium relies on communication between the myocardium and the endocardium, thereby coordinating growth of the two layers: specifically, myocardial expansion appears to relay tension to the underlying endocardium, which promotes proliferation via a VE-cadherin-Yap1 signaling pathway (Bornhorst et al., 2019). In light of these findings, it is interesting that our observations hint at coordinated expansion of the OFT myocardium and endocardium, as well as the possibility that myocardial contractility regulates OFT endocardial proliferation. It will therefore be valuable for future studies to examine Yap1 localization in the OFT endocardium, as was recently done in the OFT smooth muscle (Duchemin et al., 2019), and to assay OFT endocardial proliferation in *yap1*-deficient embryos.

Altogether, our data open new avenues for grasping the etiology of the variety of cardiac birth defects that include abnormalities in OFT morphology. Defects in early steps of OFT assembly can lead to complications in OFT remodeling and septation, which are prevalent features in congenital heart disease (Neeb et al., 2013; Pierpont et al., 2018). Given that many types of structural and physiological anomalies can lead to deficiencies in cardiac function *in utero*, our demonstration that cardiac function promotes OFT expansion by regulating both endocardial proliferation and endothelial addition provides new models to consider for the origins of multiple types of OFT defects. Taken together with our identification of a new role for Acvrl1 during OFT development, our work provides an important step forward in connecting cardiac function to specific mechanosensitive pathways and the cell behaviors that they regulate, and significantly advances our understanding of the many ways in which form follows function during organogenesis.

## MATERIALS AND METHODS

### Zebrafish

Embryos were generated by breeding wild-type zebrafish or zebrafish heterozygous for the *tnnt2a* mutant allele *sih^b109^*(Sehnert et al., 2002), the *myh7* mutant allele *haf^sk46^* (Auman et al., 2007), the *myh6* mutant allele *wea^m58^* (Berdougo et al., 2003), or the *acvrl1* mutant allele *vbg^y6^* (Roman et al., 2002), carrying the transgenes *Tg(fli1a:negfp)^y7^*(Roman et al., 2002), *Tg*(*kdrl*:*grcfp*)*^zf528^*(Cross et al., 2003), *Tg*(*kdrl*:*HsHRAS*-*mCherry*)*^s896^*(Chi et al., 2008), *Tg(kdrl:gfp)^la116^* (Choi et al., 2007), or *Tg*(*myl7*:*H2A*-*mCherry*)*^sd12^* (Schumacher et al., 2013). *klf2* mutant embryos were generated by crossing fish that were heterozygous for the mutant allele *klf2a^bns11^* and homozygous for the mutant allele *klf2b^bns12^* (Kwon et al., 2016). The transgene *Tg*(*kdrl*:*dendra*) was generated through modification of the *Tg*(*kdrl*:*HsHRAS*-*mCherry*) construct (Chi et al., 2008), followed by stable integration to create the *Tg*(*kdrl*:*dendra*)*^sd8^*line. Embryos homozygous for *sih^b109^*, *haf^sk46^*and *wea^m58^* were identified based on their contractility defects (Auman et al., 2007; Berdougo et al., 2003; Sehnert et al., 2002). Embryos homozygous for the *acvrl1* mutation *vbg^y6^* were identified based on their vascular phenotypes, including dilated cranial vessels and reduced blood flow in the tail (Roman et al., 2002). *klf2* mutants were genotyped using high-resolution melt analysis with the primers GACACCTACTGCCAACCGTCTC (F) and GGGAAAGCAGGCCTGACTAGGAAT (R) (*klf2a*) or CGTGGCACTGAACACAGAC (F) and GCTGAGATCCTCGTCATC (R) (*klf2b*) (Kwon et al., 2016; Rasouli et al., 2018).

### Injections

For knockdown experiments, embryos were injected at the 1-4 cell stage with previously characterized morpholinos: 4 ng of anti-*tnnt2a* (ZFIN: MO1-*tnnt2a*) (Sehnert et al., 2002), 2.5 ng of anti-*amhc* (ZFIN: MO1-*amhc*) (Berdougo et al., 2003), or 2.5 ng of anti-*acvrl1* (ZFIN: MO3-*acvrl1*) (Corti et al., 2011). In all cases, the morphant phenotype was assessed qualitatively for severity and was found to phenocopy the mutant phenotype. No signs of toxicity were observed.

### Immunofluorescence

Whole-mount immunofluorescence was performed using a modified version of previously described protocols (Alexander et al., 1998; Cooke et al., 2005; Zeng & Yelon, 2014). Briefly, embryos were dechorionated and then fixed in 2% methanol-free formaldehyde (Thermo Scientific, 28908) for 10 minutes, rinsed with PBS and deyolked with a solution containing 0.2% saponin (Sigma, S4521) and 0.5% Triton X-100 (G Biosciences, 786513) in PBT (PBS containing 0.1% Tween 20 (Sigma, P9416)). Embryos were then rinsed with PBT and fixed overnight with fresh 4% paraformaldehyde (PFA) (Sigma, P6148). The next day, embryos were washed with PBT, treated with 10 μg/mL proteinase K for 8 minutes (36 hpf embryos) or 13 minutes (51 hpf embryos) to facilitate antibody penetration, and fixed again for 20-30 minutes using 4% PFA. After washing with PBT to remove fixative, embryos were blocked with 2 mg/mL Bovine Serum Albumin (Sigma, A9647) and 10% goat serum in PBT overnight at 4°C. Embryos were incubated with primary antibody in block solution as follows: 1:50 dilution of mouse anti-Alcama (Developmental Studies Hybridoma Bank, Zn-8 supernatant), 1:400 dilution of rabbit or chicken anti-GFP (Life Technologies, A11122 or A10262) or 1:1000 dilution of rabbit anti-dsRed (Clontech, 632496), overnight at 4°C. Embryos were then washed extensively with PBT and incubated with the following secondary antibodies in PBT: 1:150 dilution of goat anti-mouse AlexaFluor 594 (Life Technologies, A11012) or goat anti-mouse AlexaFluor 647 (Life Technologies, A21446), 1:400 dilution of goat anti-rabbit AlexaFluor 488 (Life Technologies, A11008) or goat anti-chicken AlexaFluor 488 (Life Technologies, A11039) or goat anti-rabbit AlexaFluor 594 (Life Technologies, A11012), overnight at 4°C. TO-PRO3 (Invitrogen, T3605, 1:500) or DAPI (Invitrogen, D1306, 1:1000) were added during secondary antibody incubation as needed. After thorough washing with PBT, embryos were mounted in 50% glycerol or SlowFade Gold anti-fade reagent (Invitrogen, S36936).

### Proliferation assays

For BrdU treatments, embryos were soaked in freshly prepared 5 mg/mL BrdU in E3 medium from 36 hpf to 51 hpf, followed by fixation and detection (Dietrich et al., 2014). Our EdU incorporation assay was adapted from a previously described protocol (Hesselson et al., 2009), with slight modifications: embryos were injected pericardially or into the yolk with 1 nL EdU solution (200 μM EdU, 2% DMSO, 0.1% phenol red) at 36 hpf, followed by fixation at 51 hpf, whole-mount immunostaining as described above, and Click-iT detection as per manufacturer’s guidelines (Invitrogen, C10339). Proliferation indices were calculated using the following formula: proliferation index (%) = (number of EdU+ or BrdU+ cells/total number of cells) x 100).

### Imaging and photoconversion

Fixed samples were imaged from the left or left-ventral side on a Leica SP5 confocal laser-scanning microscope, using a 25x water objective and a slice thickness of 0.5 µm, with the exception of the experiments in Figure 3 and Figure S3-1, where the slice thickness was 0.29 µm. For live imaging, embryos at 36 or 51 hpf were anesthetized using 1 mg/mL tricaine and mounted in 1% low-melt agarose containing 1 mg/mL tricaine for imaging on a Zeiss Z.1 Light Sheet microscope. In some cases, embryos were treated with relaxation buffer for 1 hour prior to analysis, as previously described (Huang et al., 2009; Lin et al., 2012); however, comparison of treated and untreated embryos yielded no difference in OFT measurements (data not shown). 36 hpf embryos were exposed to 0.5 mg/mL tricaine for 15-30 minutes prior to full anesthesia, since this exposure leads to mild pericardial edema that improves optical access to the outflow tract. This strategy was also employed prior to photoconversion at 36 hpf, which was performed on embryos mounted in 2% methylcellulose containing 1 mg/mL tricaine, oriented with their left-ventral side towards a 25x water objective on a Leica SP5 confocal laser-scanning microscope. The slight ventral tilt allows for a clear view of the OFT and the right side of the first aortic arch (AA1), which influenced our choice to consistently perform photoconversion on the right side of the bilateral AA1 structure. A region-of-interest (ROI) was exposed to UV light at 25% laser intensity until red signal was clearly visible; this degree of UV exposure was not toxic and did not impair embryonic development. Embryos were imaged post-photoconversion with low Argon ion laser intensity (15%) to assess specificity: if regions beyond the intended ROI were photoconverted in an embryo, it was not analyzed further. Photoconverted embryos were then maintained until 51 hpf and imaged in 1 mg/mL tricaine.

### Defining landmarks in the OFT

Since there is currently no available molecular marker that is specific for the OFT at the stages analyzed, we chose to define OFT boundaries using morphological criteria, as detailed below. We used the following anatomical terminology to describe the OFT: proximal and distal ends of the OFT, with respect to the arterial pole of the ventricle, and inner and outer curvatures of the OFT, where the inner curvature (IC) is the lesser, concave surface of the curved OFT (continuous with the outer curvature of the ventricle) and the outer curvature (OC) is the greater, convex curvature of the OFT (continuous with the inner curvature of the ventricle) (Fig. 1A,B).

The proximal boundary of the OFT was defined as the cross-sectional plane at the morphological constriction of the myocardium between the ventricle and the OFT, as visualized using anti-Alcama in fixed embryos, and *Tg*(*myl7*:*H2A*-*mCherry*) in live embryos; this constriction is evident along the entire myocardial circumference. At 36 hpf, this cross-sectional plane is perpendicular to the proximal-distal axis of the OFT (Fig. 1A). By 51 hpf, due to the curvature of the OFT, this plane is typically angled such that its IC edge is tilted distally compared to its OC edge (Fig. 1B). We noted qualitative differences in cell morphologies flanking the proximal boundary of the OFT: the ventricular cardiomyocytes adjacent to the boundary appear to have a larger surface area than do the OFT cardiomyocytes that lie on the other side of the boundary.

The distal boundary of the OFT was determined primarily by the bifurcation point where the OFT endocardium meets the first aortic arch (AA1), but also considering the distal edge of Alcama localization in fixed embryos or *Tg*(*myl7*:*H2A*-*mCherry*) expression in live embryos. At 36 hpf, some Alcama is routinely found around the aortic arches in addition to around the OFT (Fig. 1A). Therefore, the OFT distal boundary is defined as the plane through the bifurcation point of AA1, proximal to the distal edge of Alcama localization. By 51 hpf, the distal end of Alcama localization usually coincides with the AA1 bifurcation point, especially on the OC side (in a typical batch of 10 embryos, 8 exhibited this characteristic) (Fig. 1B); in these cases, either location can be considered as the distal boundary of the OFT. However, in embryos where the distal edge of Alcama localization is proximal to the AA1 bifurcation point, the former is considered to be the OFT distal boundary. As with the OFT proximal plane, the distal plane is perpendicular to the proximal-distal axis of the OFT at 36 hpf, but, by 51 hpf, it is tilted such that the OFT OC ends more distally than its IC (Fig. 1A,B).

Overall, we found the proximal and distal boundaries of the OFT to be highly reproducible and stereotypical, as suggested by the low SEM for our cellular and volumetric analyses in the wild-type OFT (Figs. 1S,W and 3U,V). These boundaries were also clearly and routinely discernible in mutant and morphant OFTs, consistent with the low SEMs observed in our cellular analyses in these contexts as well (Figs. 3U,V; 4K; S4-1E; 7C; S7-1D; and S7-2C). Embryos in which the proximal or distal boundaries of the OFT were not clear, due to poor staining or imaging angle, for example, were not analyzed further.

### Morphometrics

All morphometric analyses were performed using Imaris (Bitplane) software, unless mentioned otherwise. For cell counting (Figs. 1, 2, 4, S4-1, 5, S5-1, 7, S7-1 and S7-2), the “spots” function in Imaris was used to count endocardial or myocardial nuclei in the region demarcated as the OFT. For counting cells within a middle cross-section, we selected the plane through the central points in the OC and IC along the proximal-distal axis of the OFT. In *klf2* mutants and their siblings (Fig. S7-1), due to technical difficulties associated with cytoplasmic diffusion of nuclear GFP, the nuclear marker DAPI was masked using the green channel in *Tg(fli1a:negfp)* embryos to facilitate endocardial cell counting. Similarly, since *acvrl1* mutants and wild-type siblings did not contain an endocardial nuclear marker (Fig. S7-2), the nuclear marker TO-PRO-3 was masked using the green channel in *Tg*(*kdrl*:*gfp*) embryos for endocardial cell counting.

All cell area and circularity measurements (Figs. 1 and S4-2) were performed using ImageJ (Fiji) software as described previously (Auman et al., 2007). Briefly, the OC and IC walls of the OFT were partially reconstructed (∼5 μm thick for the myocardium and ∼0.5 μm thick for the endocardium), using the slices in which lateral boundaries of cells were clearly visible, and then collapsed into maximum intensity projections. Cellular boundaries were traced using immunofluorescence for Alcama, a basolateral marker in cardiomyocytes (Jimenez-Amilburu et al., 2016), and for mCherry on endocardial cell surfaces in *Tg*(*kdrl*:*HsHRAS*-*mCherry*) embryos.

To perform OFT endocardial volumetric analysis (Figs. 3 and S3-2), a smoothed surface (surfaces detail = 1) was created automatically using cytoplasmic GFP in *Tg*(*kdrl*:*grcfp*) embryos. The surface was then cut at the proximal and distal boundaries, defined as described above. We considered the volume of this surface to be the enclosed volume, since the surface “fills” the endocardial lumen of the OFT. For cross-sectional area analysis, areas of triangles formed by three points along the circumference of the cross-section were calculated using Heron’s formula. A summation of the areas of all such triangles provided the total cross-sectional area of that plane.

To perform OFT myocardial volumetric analysis (Fig. 3), a surface was created manually using the “contour surfaces” function in Imaris, so as to obtain the enclosed volume. In multiple cross-sections through the OFT, the apical boundary of Alcama localization was drawn manually, and the three-dimensional surface was created by connecting these two-dimensional boundaries.

### Statistics and replication

Statistical analysis was performed using Prism (GraphPad). First, a Shapiro-Wilk test was run to test normality of data. If data were normally distributed, a Student’s T-test (two-tailed) was performed to compare data sets. If the data were not normally distributed, a Mann-Whitney U-test was performed to compare data sets. For all experiments, ≥2 technical replicates and ≥5 biological replicates were analyzed, unless mentioned otherwise in cases when alternative methodologies were used to complement observations. Technical replicates (N) were defined as the number of independent experiments performed, on different dates with fresh reagents. Biological replicates (n) were typically defined as the total number of embryos analyzed for each category; in experiments where cell shape or size was measured, the number of cells analyzed is also provided. Specific N and n values are presented in figure legends.

## ACKNOWLEDGEMENTS

We are grateful for helpful discussions with K. Cooper, X. Sun, J. Posakony, A. Chisholm, E.M. Collins, and members of the Yelon lab, for expert zebrafish care by T. Sanchez and H. Kwan, for reagents and advice provided by A. Anbalagan and S.J. Rasouli, and for light-sheet microscopy assistance from J. Santini and M. Erb.

## AUTHOR CONTRIBUTIONS

PS: Conceptualization, Methodology, Investigation, Formal Analysis, Writing - Original Draft Preparation, Writing - Review & Editing.

GLMB: Investigation, Resources, Writing – Review & Editing. HY: Resources, Writing – Review & Editing.

NCC: Conceptualization, Supervision, Resources, Writing – Review & Editing.

BLR: Conceptualization, Supervision, Resources, Writing – Review & Editing.

DYRS: Conceptualization, Supervision, Resources, Writing – Review & Editing.

DY: Conceptualization, Supervision, Methodology, Formal Analysis, Writing - Original Draft Preparation, Writing - Review & Editing, Project Administration, Funding Acquisition.

## FIGURE SUPPLEMENT LEGENDS

**Figure 2 – Figure Supplement 1.**
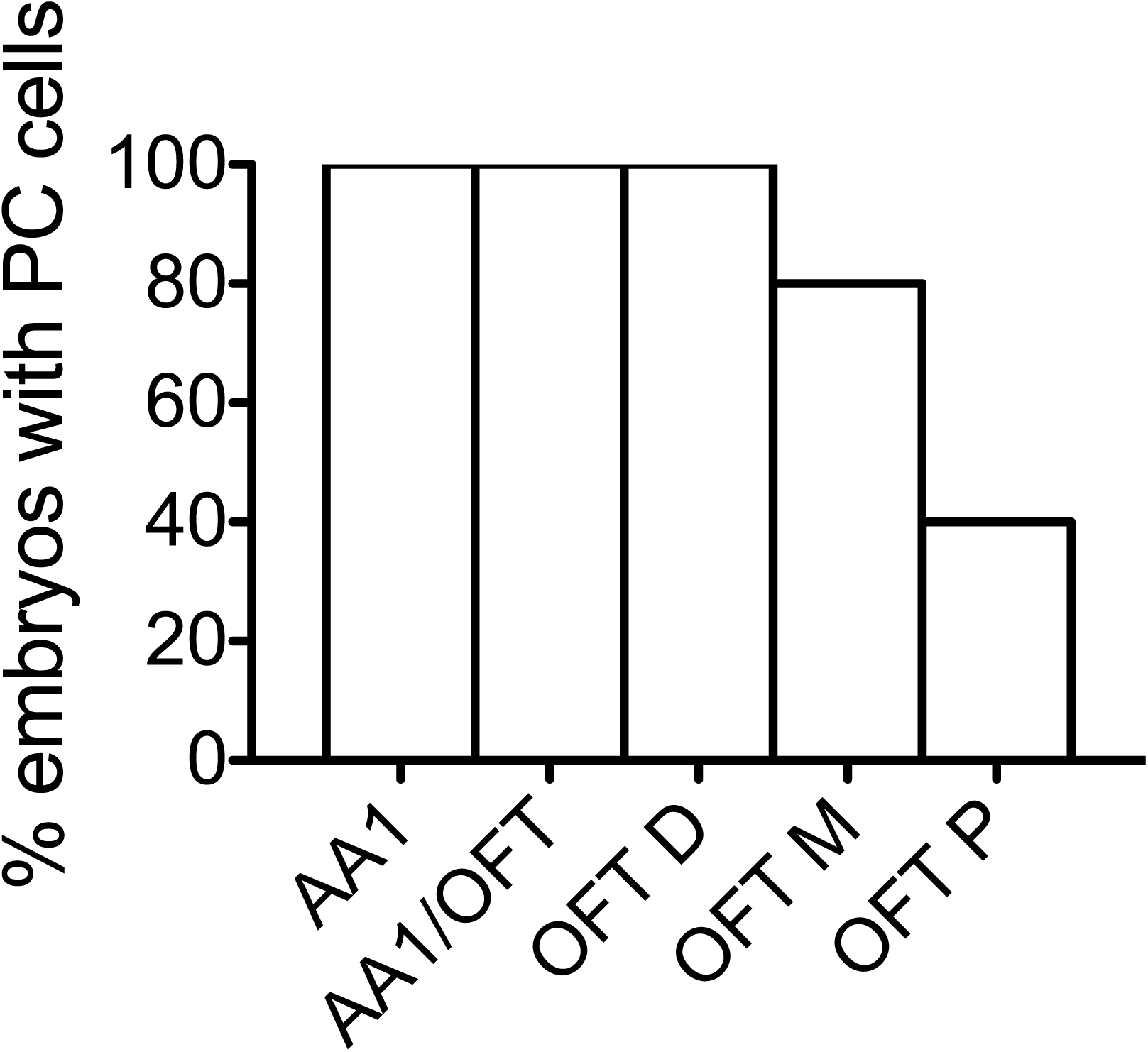
Displacement of endothelial cells from AA1 into the OFT. Bar graph indicates the percentage of embryos in which labeling of AA1 endothelium via photoconversion at 36 hpf resulted in detection of photoconverted (PC) cells in AA1, at the junction of AA1 and the OFT (AA1/OFT), in the distal third of the OFT (OFT D), in the middle third of the OFT (OFT M), or in the proximal third of the OFT (OFT P) at 51 hpf (n=5; N=2). In the particular example shown (Fig. 2T-V), labeled cells were found in AA1, AA1/OFT, OFT D, and OFT M, but not in OFT P.

**Figure 3 – Figure Supplement 1.**
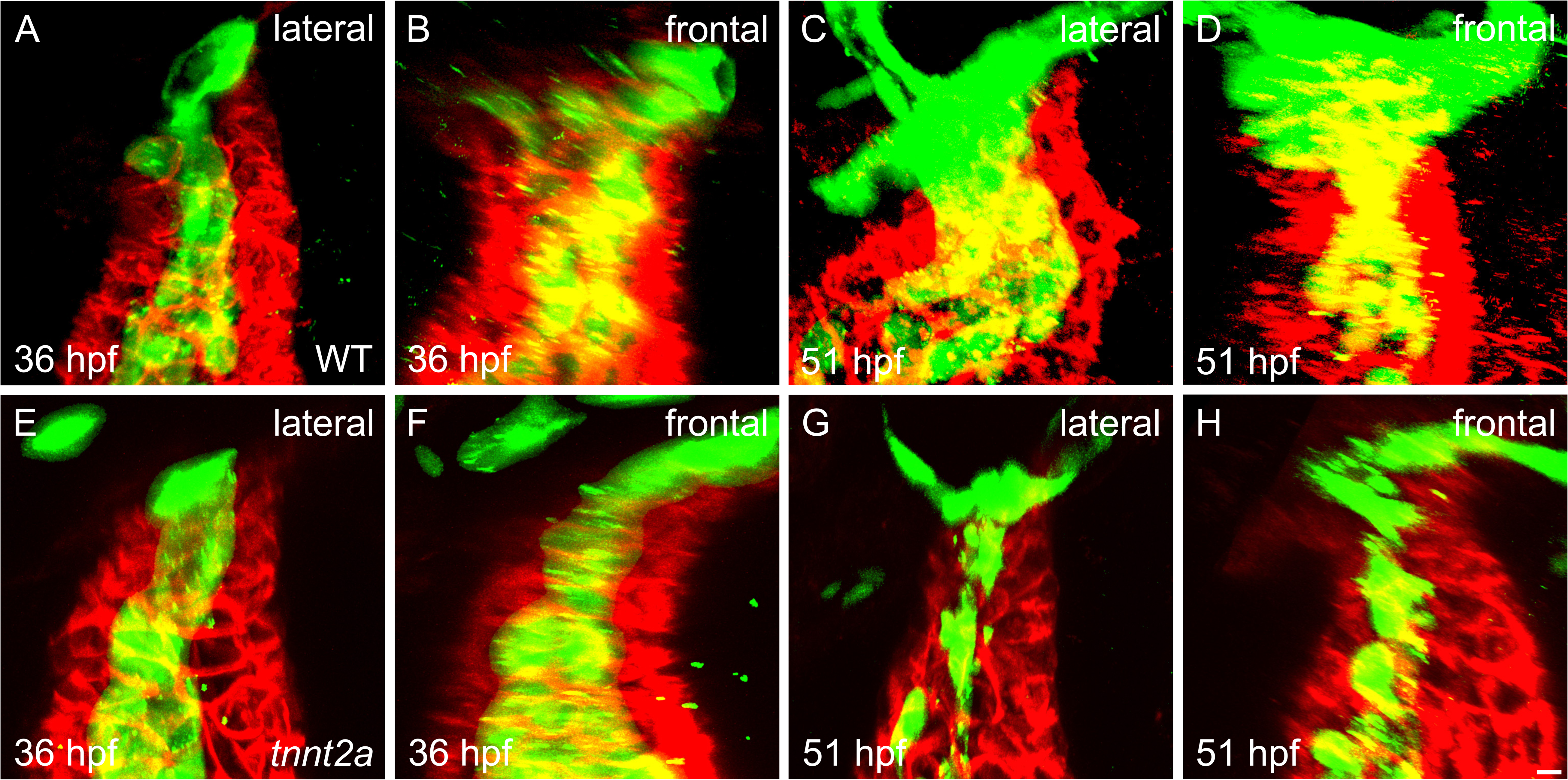
Cardiac function influences OFT endocardial growth. **(A-H)** Three-dimensional reconstructions of lateral (A,C,E,G) and frontal (B,D,F,H) views of the OFT in WT (A-D) and *tnnt2a* mutant (E-H) embryos, labeled by immunofluorescence for Alcama (red) and *Tg*(*kdrl*:*grcfp*) (green), the surface representations of which are displayed in Fig. 3B,C,E,G,H,J,L,M,O,Q,R,T. Confocal Z-slices were captured in the lateral orientation at low sampling intervals and rotated, using Imaris software, to generate frontal views. Scale bar: 5 µm.

**Figure 3 – Figure Supplement 2.**
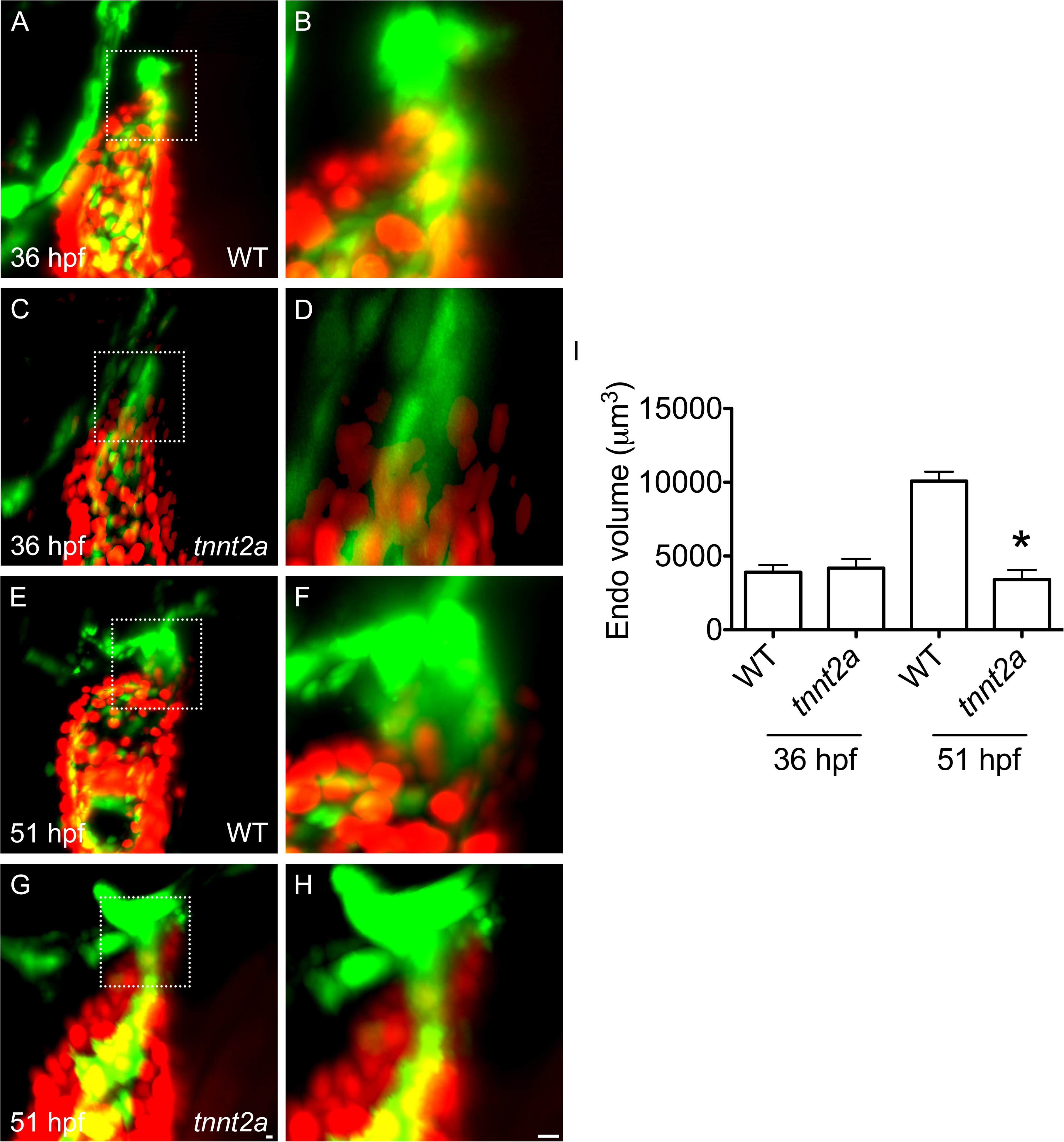
Live imaging confirms that OFT endocardial growth depends upon cardiac function. **(A-H)** Three-dimensional reconstructions of lateral views of live embryos carrying the transgenes *Tg*(*kdrl*:*grcfp*) and *Tg*(*myl7*:*H2A*-*mCherry*), in which the endocardium is labeled with cytoplasmic GFP (green) and the myocardial nuclei are labeled with mCherry (red). Views show the OFT on top of the ventricle (A,C,E,G), as well as a closer view of the OFT (B,D,F,H; correspond to boxed regions in A,C,E,G). At 36 hpf, OFT morphology appears similar in WT (A,B) and *tnnt2a* mutant (C,D) embryos. However, by 51 hpf, the volume of the *tnnt2a* mutant OFT endocardium (G,H) is significantly smaller than in WT (E,F), as quantified in (I) (asterisk indicates significant difference from WT at same stage, p<0.0001, Student’s T test). **(I)** Bar graph indicates mean and SEM (n=7,3 at 36 hpf; N=1; n=6,5 at 51 hpf; N=2). While the mean endocardial volume for each set of live embryos is higher than its counterpart from fixed embryos (Fig. 3U), the degree of difference in endocardial volume between 36 hpf and 51 hpf WT embryos (p<0.0001, Student’s T test), as well as between WT and *tnnt2a* mutants at 51 hpf, is relatively similar between live and fixed samples. Scale bars: 5 µm.

**Figure 4 – Figure Supplement 1.**
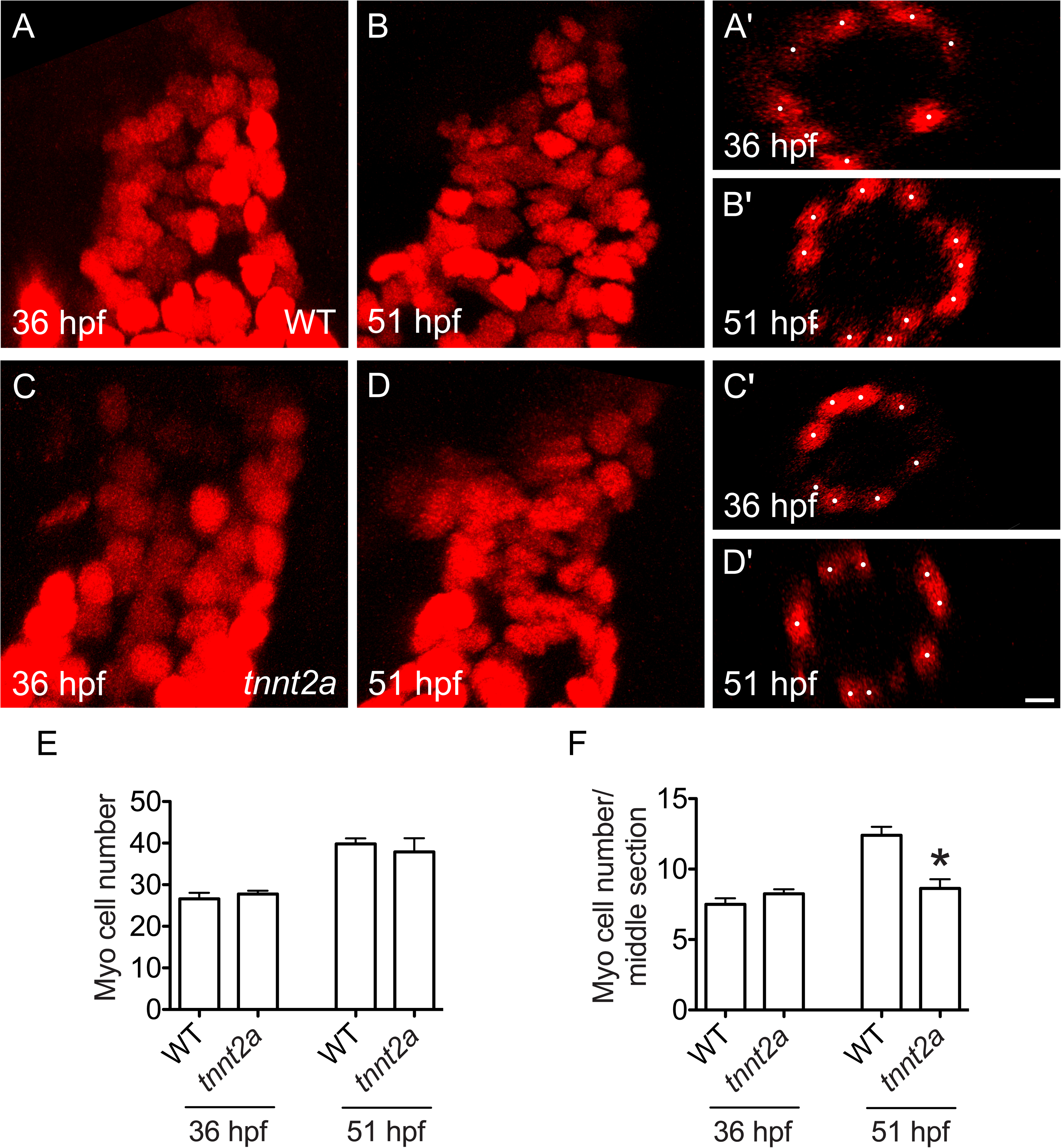
Cardiac function influences myocardial arrangement in the OFT. **(A-D)** Three-dimensional reconstructions (A-D) and middle cross-sections (A’-D’) of the OFT (as in Fig. 1K,L) illustrate the number of OFT myocardial cells at 36 (A,C) and 51 hpf (B,D). At both stages, the total number of myocardial cells in the OFT is similar in WT (A,B) and *tnnt2a* mutant embryos (C,D), as quantified in (E). However, whereas the number of cells in a middle section is comparable between WT (A′) and *tnnt2a* mutants (C′) at 36 hpf, it is significantly reduced in *tnnt2a* mutants by 51 hpf (D′) compared to WT (B′), as quantified in (F). **(E,F)** Bar graphs indicate mean and SEM (n=6,8 for 36 hpf; 5,8 for 51 hpf; N=2; asterisk indicates significant difference from WT at same stage; p<0.05, Student’s T test). Scale bar: 5 µm.

**Figure 4 – Figure Supplement 2.**
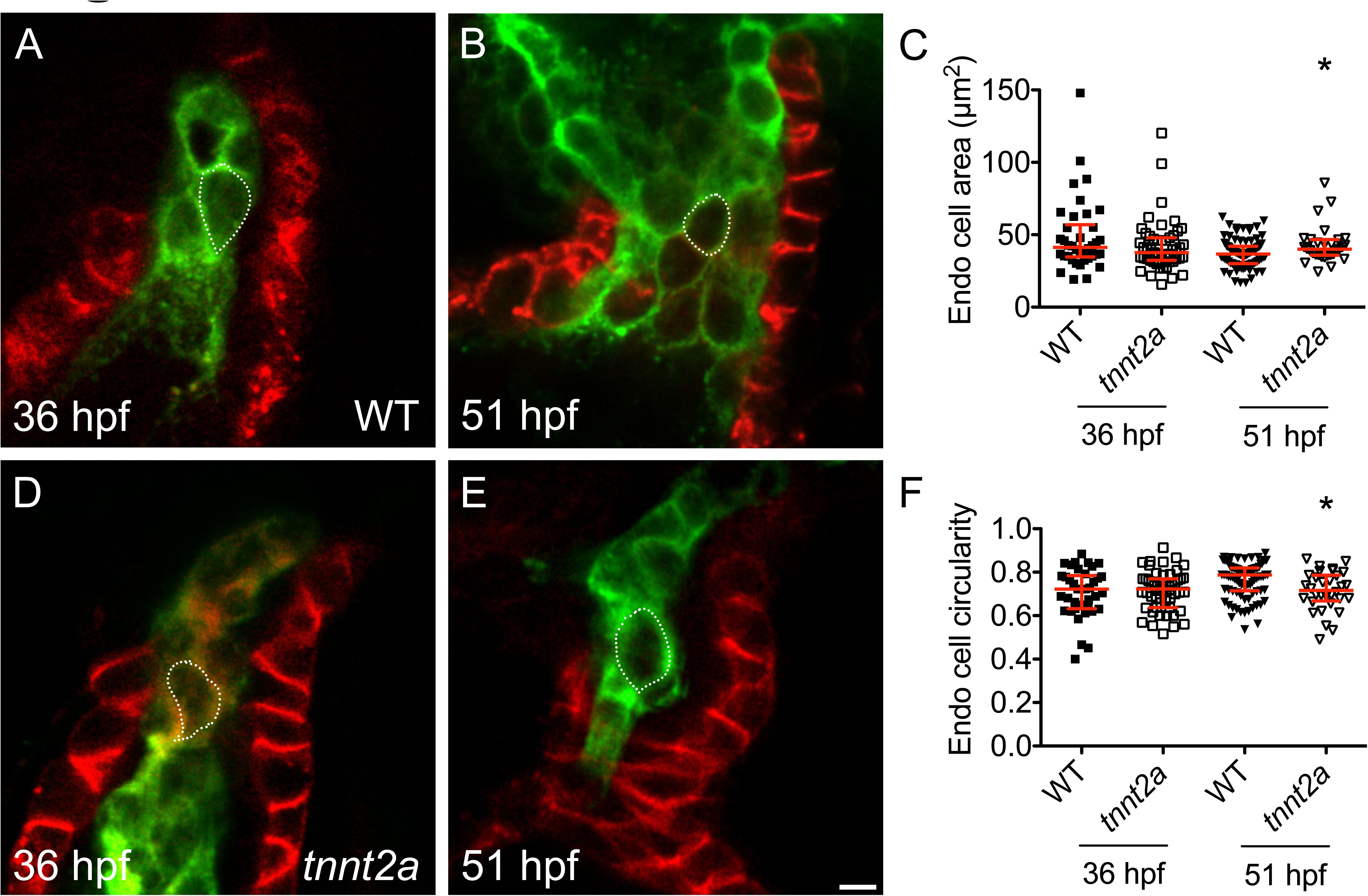
Cardiac function influences endocardial cell size and shape in the OFT. **(A,B,D,E)** Partial reconstructions illustrate representative OFT endocardial walls (as in Fig. 1E,F,IJ). At 36 hpf, OFT endocardial cells are similar in area and circularity in WT (A) and *tnnt2a* mutant (D) embryos, as quantified in (C,F). However, by 51 hpf, whereas OFT endocardial cells become smaller (C, p<0.05, Mann-Whitney test) and rounder (F, p<0.005, Mann-Whitney U test) in WT (B), they remain relatively large and elongated in *tnnt2a* mutants (E). In these data sets, OC and IC cells are grouped together, due to technical challenges associated with distinguishing the walls within the collapsed endocardium in *tnnt2a* mutants at 51 hpf. **(C,F)** Red lines represent median and interquartile range (n=9, 8, 9, 5 embryos; 38, 51, 87, 30 cells; N= 3; asterisks indicate significant difference from WT at same stage, p<0.05, Mann-Whitney U test). Scale bar: 5 µm.

**Figure 5 – Figure Supplement 1.**
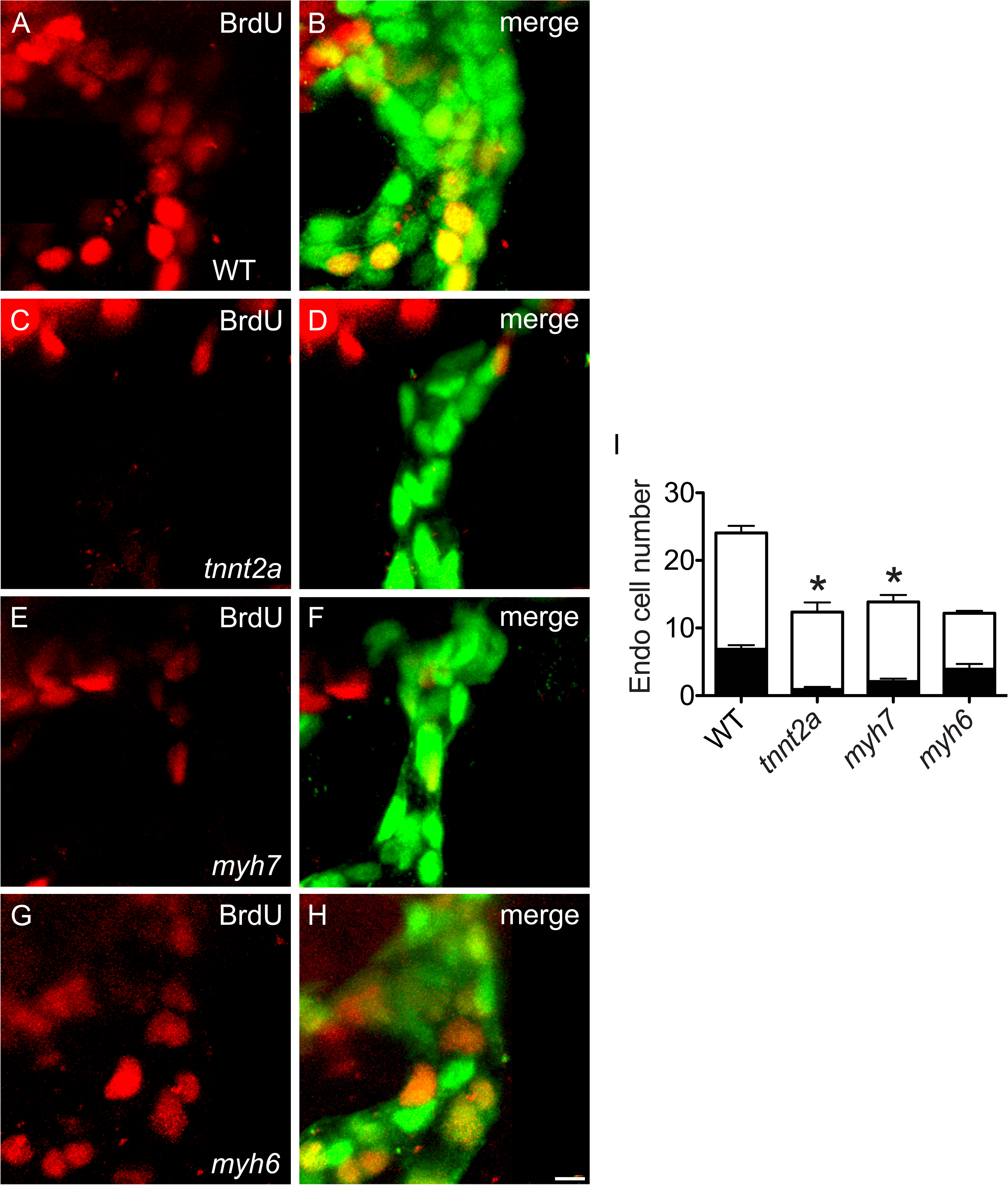
BrdU incorporation assay confirms reduction of OFT endocardial proliferation in *tnnt2a* and *myh7* mutant embryos. **(A-H)** Three-dimensional reconstructions of the OFT (as in Fig. 2F,G) illustrate BrdU incorporation between 36 and 51 hpf in WT (A,B), *tnnt2a* mutant (C,D), *myh7* mutant (E,F) and *myh6* mutant (G,H) embryos. **(I)** Black and white bars in graph represent mean number of BrdU+ and BrdU- endocardial cells, respectively, with SEM. BrdU incorporation is significantly reduced in *tnnt2a* mutants (PI=8.2 ± 3.8%) and *myh7* mutants (PI=13.8 ± 2.7%), but not in *myh6* mutants (PI=29.1 ± 4.4%), compared to WT (PI=28.5 ± 2.2%) (n=21, 12, 12, 12; N=2; asterisks indicate significant differences in PI compared to WT; p<0.001, Student’s T test). Scale bar: 5 µm.

**Figure 6 – Figure Supplement 1.**
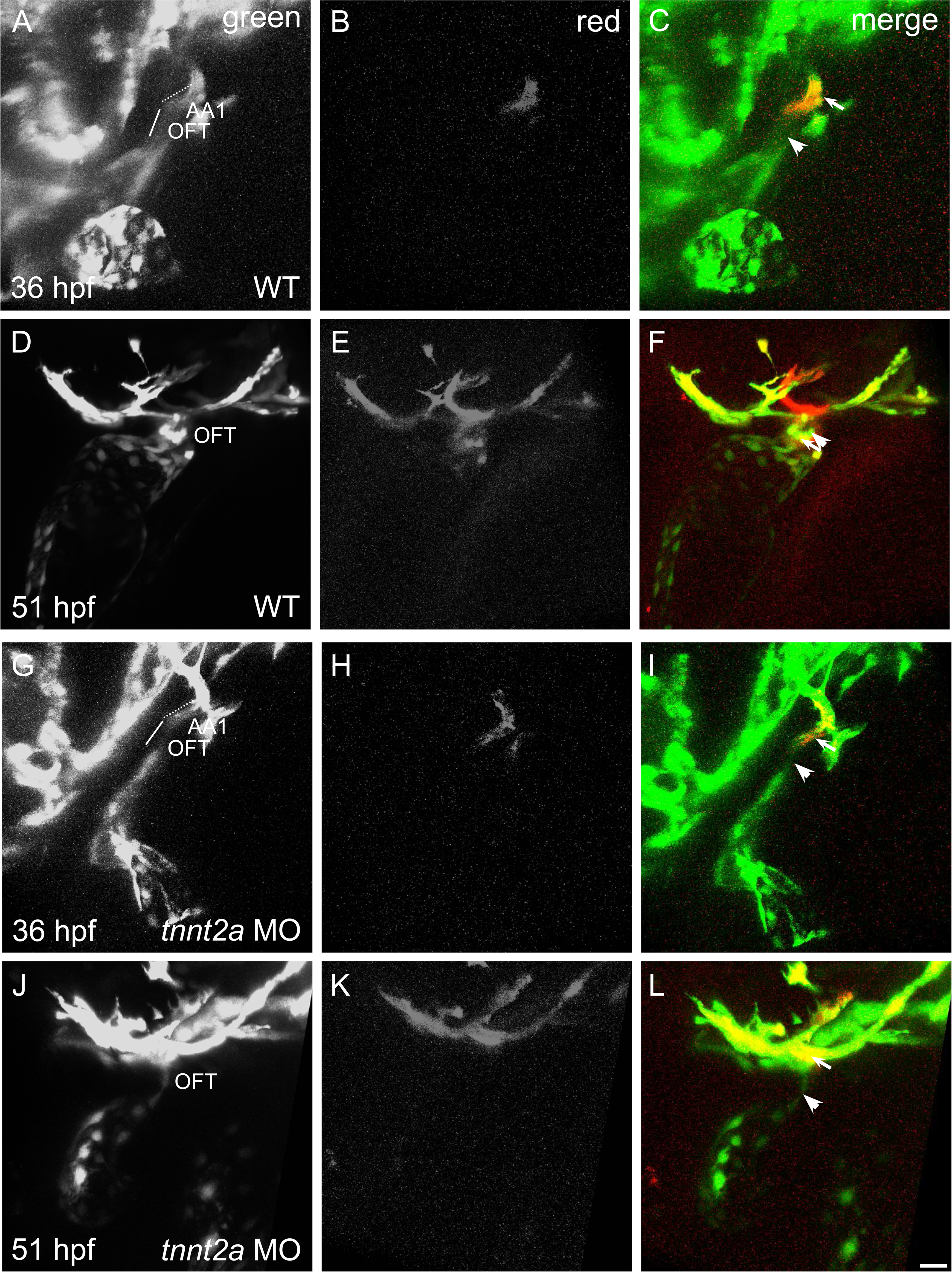
*tnnt2a* morphants recapitulate OFT endothelial addition defects observed in *tnnt2a* mutants. **(A-F)** Three-dimensional reconstructions (as in Fig. 2Q-V) demonstrate that photoconversion of a portion of AA1 endothelium in WT at 36 hpf (A-C) results in the presence of labeled cells in the OFT by 51 hpf (D-F) (n=5; N=2). **(G-L)** This is not the case in *tnnt2a* morphants, which lack labeled cells in the OFT at 51 hpf (J-L) following photoconversion within the AA1 endothelium at 36 hpf (G-I) (n=5; N=2). Arrowheads indicate examples of cells that were not labeled by photoconversion (green only), and arrows indicate examples of cells that were labeled by photoconversion (red and green). Scale bar: 20 µm.

**Figure 7 – Figure Supplement 1.**
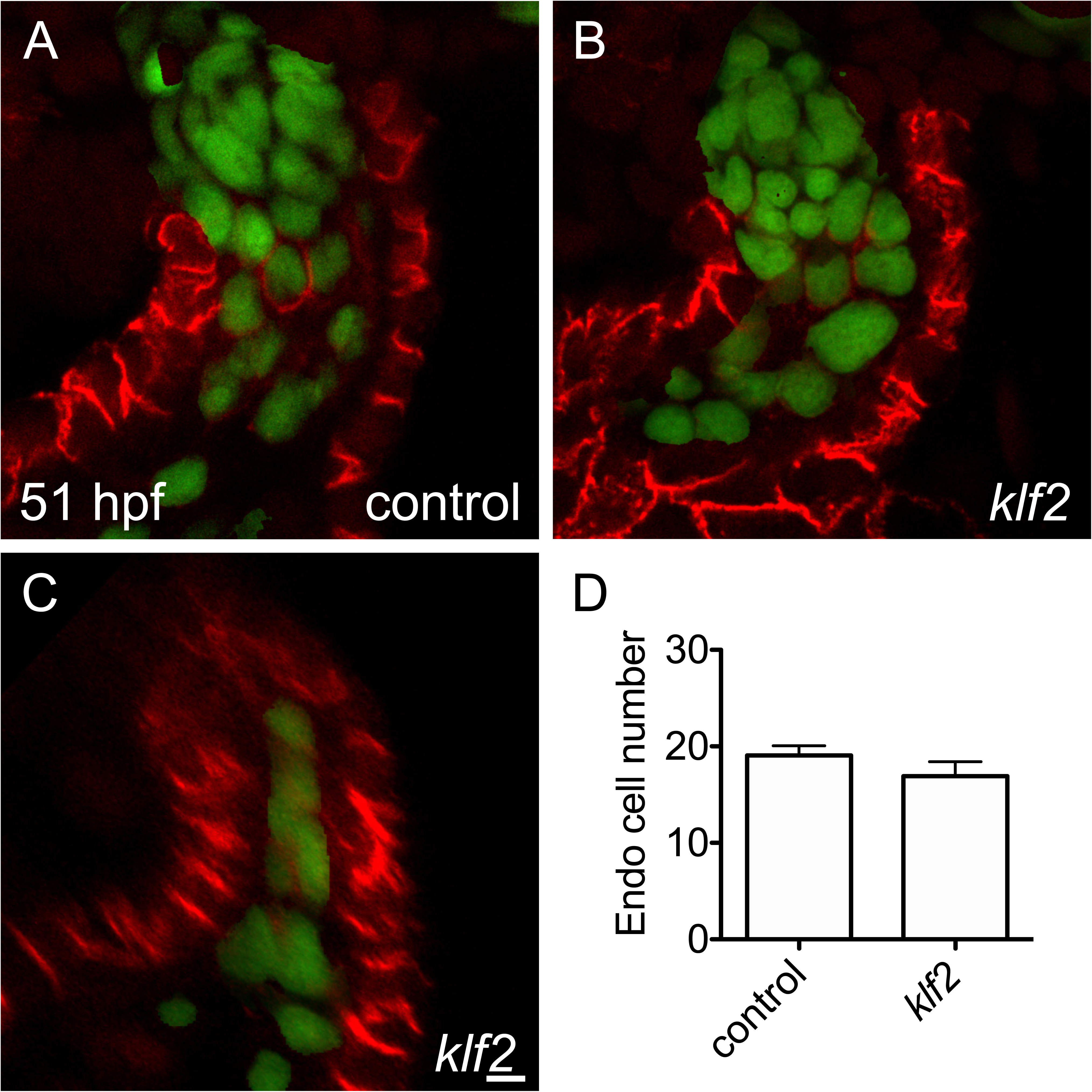
Klf2a and Klf2b are not essential for OFT endocardial expansion. **(A-C)** Immunofluorescence indicates localization of Alcama (red) in myocardial cells and expression of *Tg(fli1a:negfp)* in endocardial cells; in these images, green signal indicates the masking of the nuclear marker DAPI using GFP fluorescence (see Materials and Methods). Lateral slices through the OFT demonstrate that, at 51 hpf, in comparison to the OFT in sibling controls (A), the *klf2* mutant OFT typically exhibits a normal number of endocardial cells and a normal morphology (B; 10/11 *klf2* mutants) but can exhibit fewer endocardial cells and a collapsed morphology (C; 1/11 *klf2* mutants). **(D)** Quantitative analysis of our entire data set did not reveal a significant difference between the number of OFT endocardial cells in *klf2* mutant embryos and their sibling controls. Bar graph indicates mean and SEM (n= 18, 11; N=3). Scale bar: 5 µm.

**Figure 7 – Figure Supplement 2.**
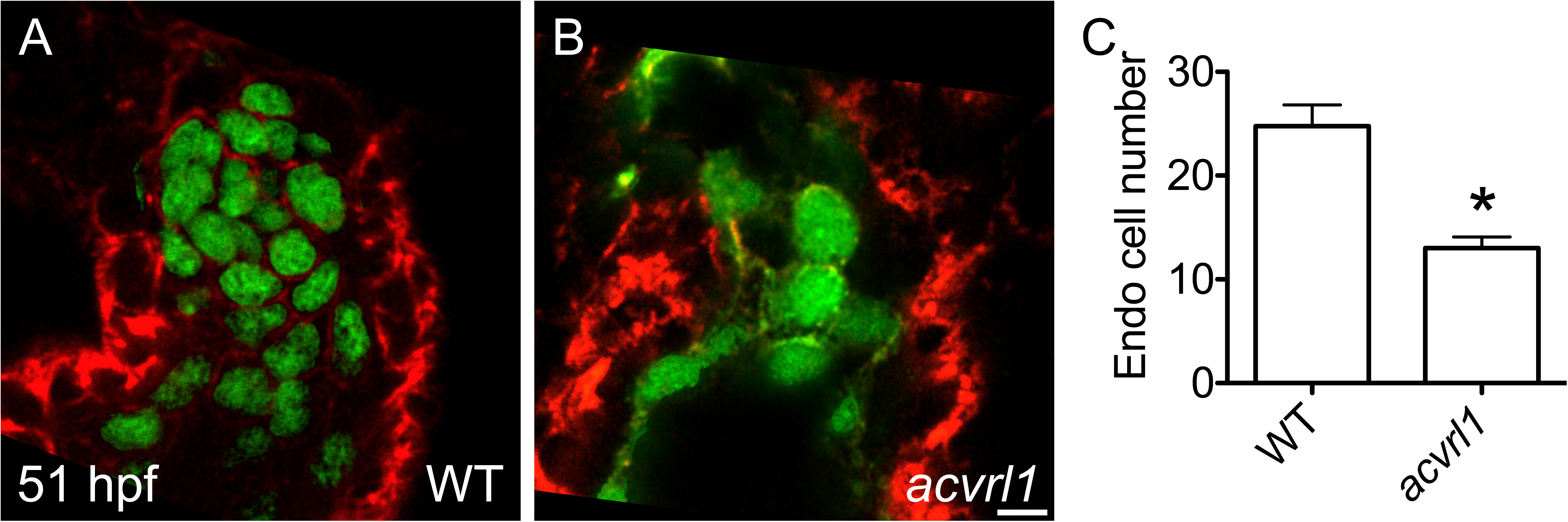
*acvrl1* mutants phenocopy *acvrl1* morphants with respect to OFT endocardial cell number. **(A,B)** Lateral slices through the OFT in WT (A) and *acvrl1* mutant (B) embryos illustrate expression of the endothelial reporter transgene *Tg(kdrl:GFP)* and labeling of the myocardium with phalloidin (red); in these images, green signal indicates the masking of the nuclear marker TO-PRO-3 using GFP fluorescence (see Materials and Methods). **(C)** Bar graph indicates mean and SEM and demonstrates that *acvrl1* mutant embryos, similar to *acvrl1* morphants, have a significant reduction in the number of OFT endocardial cells (n=4, 4; N=1; asterisk indicates significant difference from WT, p<0.01, Student’s T test). Scale bar: 5 µm.

**Figure 7 – Figure Supplement 3.**
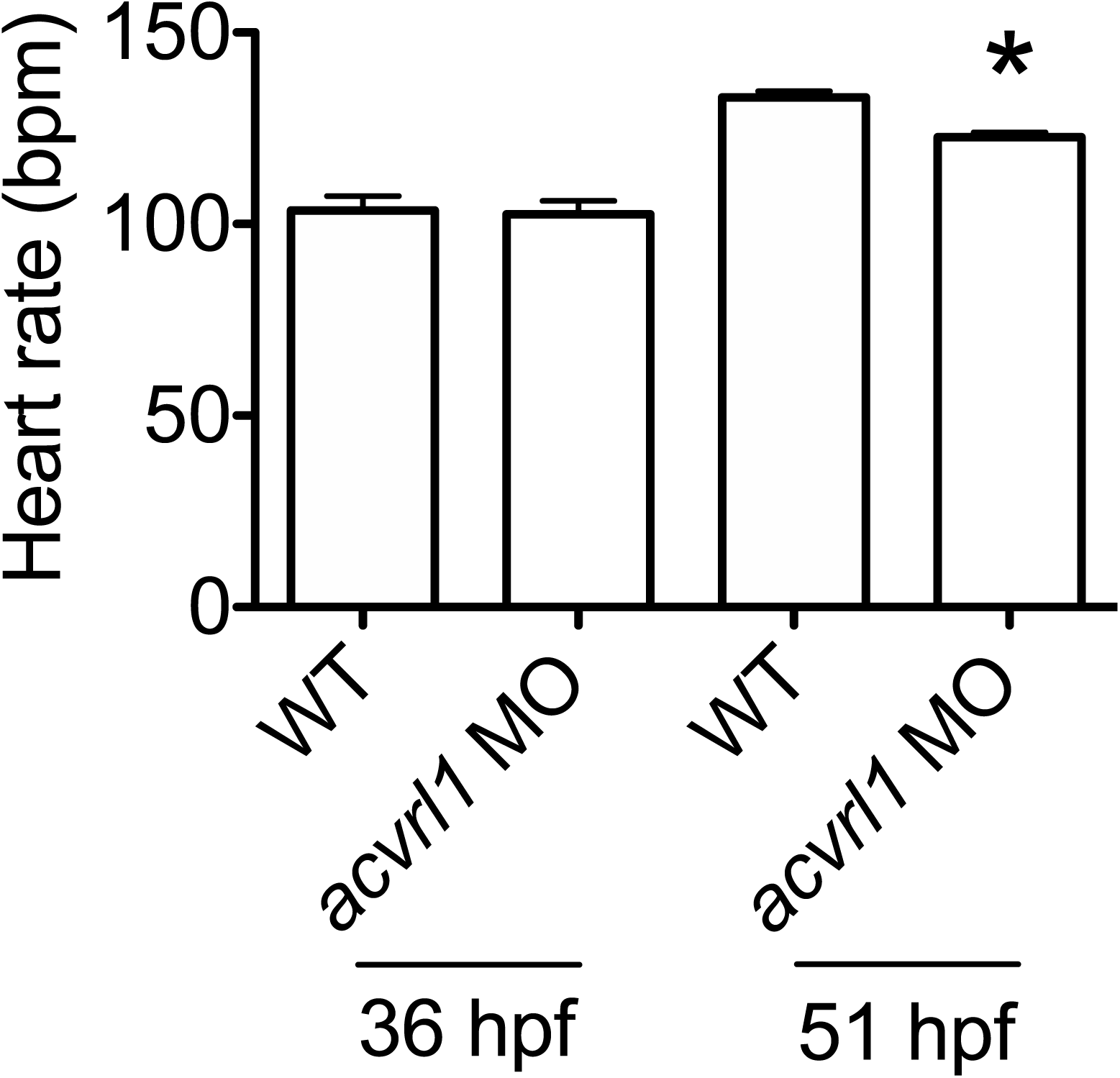
*acvrl1* morphants exhibit a reduced heart rate by 51 hpf. Heart rate was calculated by counting ventricular beats for 30 seconds under a stereoscope. At 36 hpf, the heart rate (in beats per minute (bpm)) is comparable in WT embryos and *acvrl1* morphants. However, by 51 hpf, *acvrl1* morphants exhibit a small but significant reduction in heart rate. Bar graph indicates mean and SEM (n= 4, 4, 11, 11; N=3; asterisk indicates significant difference from WT at same stage, p<0.0001, Student’s T test).

## SUPPLEMENTARY VIDEO LEGENDS

**Supplementary Video 1. Morphology of the WT OFT at 36 hpf.**

Movie illustrates a three-dimensional reconstruction of the WT OFT and ventricle at 36 hpf, as shown in Fig. 3A, beginning with a lateral view of the OFT and ventricle, zooming in on the OFT, rotating to show a ventral view, and finally transitioning to a different lateral view. Scale bars adjust to perspective. The endocardium exhibits relatively uniform width along the proximal-distal axis of the OFT.

**Supplementary Video 2. Expansion of the WT OFT is evident by 51 hpf.**

Movie (as in Video S1) illustrates a three-dimensional reconstruction of the WT OFT and ventricle at 51 hpf, as shown in Fig. 3K. The volume of the OFT endocardium and myocardium appear increased at 51 hpf, compared to their appearance at 36 hpf. The distal expansion of the OFT endocardium is especially evident in a frontal view.

**Supplementary Video 3. The *tnnt2a* mutant OFT resembles the WT OFT at 36 hpf.**

Movie (as in Video S1) illustrates a three-dimensional reconstruction of the *tnnt2a* mutant OFT and ventricle at 36 hpf, as shown in Fig. 3F. While signs of endocardial defects are becoming evident in *tnnt2a* mutants at 36 hpf, the overall morphology of the OFT endocardium still appears similar to WT at this stage.

**Supplementary Video 4. The *tnnt2a* mutant OFT endocardium collapses by 51 hpf.**

Movie (as in Video S1) illustrates a three-dimensional reconstruction of the *tnnt2a* mutant OFT and ventricle at 51 hpf, as shown in Fig. 3P. While the *tnnt2a* mutant OFT myocardium appears similar in volume to WT at 51 hpf, the *tnnt2a* mutant OFT endocardium appears as a single row of cells with no evident lumen at this stage.

**Supplementary Video 5. Endothelial cells from AA1 enter the WT OFT endocardium by 51 hpf.**

Movie depicts surfaces of areas labeled by photoconversion (red) amongst surfaces of unlabeled areas (green) in a WT embryo at 51 hpf, following labeling of a portion of the AA1 endothelium at 36 hpf, as shown in Fig. 6F. Surfaces for the red channel were generated using a lower threshold so that they overlay surfaces for the green channel. Scale bars adjust to perspective. Labeled cells within the OFT appear relatively continuous with labeled cells in AA1.

**Supplementary Video 6. Cardiac function promotes addition of endothelial cells from AA1 to the OFT endocardium.**

Movie (as in Video S5) depicts location of cells labeled by photoconversion (red) amongst unlabeled cells (green) in a *tnnt2a* mutant embryo at 51 hpf, following labeling of a portion of the AA1 endothelium at 36 hpf, as shown in Fig. 6L. While labeled cells are visible in AA1, they are absent from the OFT.

**Supplementary Video 7. Atrial function promotes endothelial displacement into the OFT endocardium.**

Movie (as in Video S5) depicts location of cells labeled by photoconversion (red) amongst unlabeled cells (green) in a *myh6* morphant embryo at 51 hpf, following labeling of a portion of the AA1 endothelium at 36 hpf, as shown in Fig. 6R. Labeled cells in the AA1 endothelium appear relatively continuous with labeled cells at the distal tip of the OFT.

**Supplementary Video 8. Acvrl1 drives endothelial addition from AA1 to the OFT endocardium.**

Movie (as in Video S5) depicts location of cells labeled by photoconversion (red) amongst unlabeled cells (green) in an *acvrl1* morphant embryo at 51 hpf, following labeling of a portion of the AA1 endothelium at 36 hpf, as shown in Fig. 8L. The absence of labeled cells in the OFT is evident despite the aberrant morphology of the OFT and heart.

